# The Olfactory Epithelium: A Critical Gateway for Pathological Tau Propagation and a Target for Mitigating Tauopathy in the Central Nervous System

**DOI:** 10.1101/2025.03.03.637437

**Authors:** Marion Dourte, Esther Paître, Mongia Bouchoucha, Emilien Boyer, Sandra O. Tomé, Emilie Doeraene, Caroline Huart, Karelle Leroy, Dietmar Rudolf Thal, Anabelle Decottignies, Bernard Hanseeuw, Nuria Suelves, Pascal Kienlen-Campard

**Affiliations:** Aging and Dementia group, Cellular and Molecular Division (CEMO), Institute of Neuroscience (IoNS), UCLouvain, Brussels, Belgium; Laboratory for Neuropathology, Department of Imaging and Pathology, Leuven Brain Institute (LBI), KU Leuven, Leuven, Belgium; Alzheimer and Other Tauopathies Research Group, ULB Neuroscience Institute (UNI), ULB Center for Diabetes Research (UCDR), Faculty of Medicine, Université Libre de Bruxelles, Brussels, Belgium; OlfactIoNS group, Institute of Neuroscience (IoNS), UCLouvain, Brussels, Belgium; Department of Otorhinolaryngology, Head and Neck Surgery, Cliniques Universitaires Saint-Luc, Brussels, Belgium; Department of Pathology, University Hospital Leuven, Leuven, Belgium; Genetic and Epigenetic Alterations of Genomes Unit, de Duve Institute, UCLouvain, Brussels, Belgium; Louvain Aging Lab, Institute of Neuroscience (IoNS), UCLouvain, Brussels, Belgium; Neurology Department, Cliniques Universitaires Saint-Luc, Brussels, Belgium

**Keywords:** Tau pathology, Olfactory system, Tau spreading, Olfaction

## Abstract

Olfactory impairment is a recognized early indicator of neurodegenerative diseases (NDs), such as Alzheimer’s disease (AD). Intracellular aggregates of hyperphosphorylated tau protein, referred to as neurofibrillary tangles (NFTs), are a hallmark of AD. NFTs are found in the olfactory bulb (OB) and entorhinal cortex (EC), both crucial for processing olfactory information. We explored the hypothesis that typical tau lesions could appear early and progress along olfactory regions to reach connected areas critically affected in AD (e.g. EC and hippocampal formation). To that end, we used transgenic PS19 mice expressing mutated human tau protein (1N4R isoform, P301S mutation). They recapitulate major phenotypes of AD, such as accumulation of NFTs, synaptic dysfunction, cognitive impairment, and neuronal loss. The presence of pathological hyperphosphorylated human tau protein (pTau) was monitored in olfactory regions: olfactory epithelium (OE), OB, piriform cortex (PC), and in connected regions of the hippocampal formation (hippocampus and EC). pTau was detected in the OE’s middle stratum and in the OB’s olfactory nerve layer (ONL) at 1.5 months. At 6 months of age, tau accumulations were found in the PC and EC, along with the CA3 region and dentate gyrus of the hippocampus. We found that olfactory function remained unaffected in PS19 mice, despite the presence of tau pathology in key regions of the olfactory system. Complete stripping of the OE by intranasal administration of ZnSO_4_ led to a significant reduction in pretangle-like tau pathology within the PC, amygdala, and EC of 6-month-old PS19 mice. Finally, we observed in human post-mortem samples that pTau signal was present in the olfactory regions (OE and OB) of patients at early Braak stages (I/II). Based on these observations, we propose that pTau could appear, due to ageing or environmental agents, in the OE and subsequently spread in a prion-like manner to the hippocampal formation along neuroanatomical connections. These findings also indicate the interest of the OE as a target for intervention aimed at mitigating the progression of tauopathy in the CNS.

## Introduction

Olfactory function deteriorates with age, likely due to a combination of peripheral and central alterations. One contributing factor is the gradual replacement of the olfactory epithelium (OE)—which resides in the nasal cavities and houses olfactory sensory neurons (OSNs)—by respiratory epithelium [12, 33]. Olfactory dysfunction is a common and early symptom of many neurodegenerative diseases (NDs), including Parkinson’s disease (PD), Alzheimer’s disease (AD) and frontotemporal lobe degeneration (FTLD) [6, 33]. FTLD and AD are primary and secondary tauopathies – characterized by abnormal tau protein aggregation and deposition in the brain [47].

Tau is predominantly found in neuronal axons associated with microtubules. It is known to play a crucial role in microtubule dynamics and stabilization to preserve the integrity of the cell’s cytoskeleton [16]. In NDs, tau hyperphosphorylation impairs its ability to associate with microtubules. Hyperphosphorylated tau proteins form the core of neurofibrillary tangles (NFTs), which are pathological hallmarks of various NDs, including AD [27]. The gradual deposition of hyperphosphorylated tau aggregates within specific areas is instrumental to AD pathology. The distribution pattern of tau lesions progresses according to a sequence known as Braak staging [10, 11]. In Braak stages I/II, NFTs are confined mainly to the (trans-)entorhinal region in the temporal lobe. Stages II/III involve limbic regions such as the hippocampal formation and parahippocampal areas, and extensive neocortical involvement is observed in the later stages IV [10, 11].

Several studies have demonstrated that approximately 90% of AD patients suffer from olfactory dysfunction in the early stages of the disease, characterized by decreased odor discrimination and identification, increased olfactory threshold and olfactory memory loss [4, 7, 69]. Olfactory dysfunction is thus considered a prodromal symptom of AD, preceding cognitive impairment [39, 57, 66]. In addition, *postmortem* analysis of olfactory regions in AD patients showed the presence of tau pathology in the olfactory bulb (OB), anterior nucleus, and piriform cortex (PC) [55, 66]. Olfactory dysfunction and the accumulation of tau pathology in the OB progressively worsen with the severity of Alzheimer’s disease [6]. Importantly, tau pathology is observed in the olfactory system in all definite AD cases and in more than 1/3 of elderly individuals with or without mild cognitive impairment associated with Braak stage II [5].

Since the olfactory system is highly conserved across species [61, 69], mouse models are relevant tools to investigate how impairment of the olfactory system could relate to typical features of human NDs. PS19 mice, also known as Tau P301S mice, were first described as a transgenic tauopathy mouse model by Yoshiyama *et al.* in 2007 [67]. They express mutant human microtubule-associated protein tau (MAPT) under the control of the mouse prion protein promoter (*Prnp*). The transgene encodes the 1N4R isoform harboring the disease-associated P301S mutation [67]. Progressive accumulation of NFTs has been observed in the neocortex, amygdala, hippocampus, brainstem, and spinal cord of PS19 mice from 6 months of age. Neurodegeneration starts around 9 months, particularly in the hippocampus [67]. Previous studies also showed that PS19 mice exhibited progressive neuronal cell loss in the OB and PC, along with increased latency in finding buried food [69].

The OE is the main olfactory receptor structure. It is present in the nasal cavity and is formed of three different cell types: OSNs, supporting cells and basal stem cells [8, 30]. The stem cells are divided into globose basal cells (GBCs), which serve as active progenitors for OSN regeneration throughout life [8, 12, 60], and horizontal basal cells (HBCs), which function as reserve stem cells [8, 30]. The somas of the OSNs are located at an intermediate depth within the OE, while their axons converge to form the axon bundles (AB). These bundles cross the cribriform plate and constitute the olfactory nerve that reaches the outer layer of the OB [12]. The respiratory epithelium, which lacks neurons, is present in the nasal cavities along with the OE. Human olfactory mucosa biopsy showed that aging leads to the replacement of the OE by respiratory epithelium [12].

In mammals, the OB structure is characterized by six cell layers, arranged from the periphery to the center: the olfactory nerve layer (ONL), the glomerular layer (GL), the external plexiform layer (EPL), the mitral cell layer (MCL), the internal plexiform layer (IPL) and the granular cell layer (GCL) [56]. The ONL contains the projections of OSNs, which form synapses with glomerular cells (mitral cells, tufted cells and periglomerular cells) in the GL. The mitral and tufted cells, which are the main output neurons of the OB, are involved in the transmission of olfactory information to other regions of the central nervous system (CNS) including the PC [56]. The PC is part of the olfactory cortex and is known to play a critical role in odor encoding and memory [69]. Projections of the OB reach the amygdala via the PC or directly to specific amygdala subregions via monosynaptic projections [45]. Other projections from the OB also reach the EC which is involved in both olfactory learning and memory [69]. The EC sends projections to the dentate gyrus and the CA3 region of the hippocampus, via the perforant path, which originates from layer II of the EC. Neurons from the dentate gyrus (DG) also project to the CA3 region via mossy fibers. Finally, neurons from the CA3 region project to the CA1 region via Schaffer collaterals [59]. This suggests that the progression of pathological forms of tau along the olfactory system could contribute to the appearance of lesions in regions particularly affected by tauopathy in AD (e.g. EC) at the earliest stages.

In this study, we used the PS19 model to monitor the progression of tau pathology across key regions of the olfactory and central nervous systems, including the OE, OB, PC, EC, and hippocampus. We examined the spread of tau pathology within these regions and evaluated its impact on olfactory function. Our findings revealed a distinct pattern of tau propagation in the olfactory system, with aberrant tau phosphorylation first detected in the OE, OB, and EC at 1.5 months. This pathology later appeared in the PC at 3 months and typical NFT-like tau accumulations were observed in both the PC, and the EC from 6 months onward. These finding parallels tau pathology observed in the olfactory system of patients with AD [6, 7]. We then investigated whether zinc sulfate (ZnSO_4_) intranasal treatment, aimed at removing accumulated pathological tau in the OE, could mitigate tau pathology in vulnerable regions of the CNS. Irrigation of the olfactory organ with ZnSO_4_ triggers the degeneration of OSNs, although the tissue regenerates within a month [32, 43]. Mice repeatedly treated with intranasal irrigation of ZnSO_4_ at 1.5, 2.5, and 4 months showed a significant decrease in NFT-like tau accumulations in the PC, EC, and amygdala at 6 months. This striking observation indicates that the accumulation of pathological tau in the OE could play a causal role in the progression of tau pathology observed in regions of the CNS involved in the processing of olfactory information, and particularly the EC and amygdala, which are affected in the early/intermediate stages of AD [11, 15, 50].

## Materials and methods

### Animals

PS19 mice were obtained from the Jackson Laboratory (B6; C3-Tg(Prnp-MAPT*P301S)PS19Vle/J; Strain #008169) and maintained on a C57BL/6 genetic background. Wild-type (WT) and heterozygous PS19 mice were crossed to obtain WT and heterozygous PS19 mice at 1.5, 3, 6 and 9 months of age. The genotype was determined by PCR analysis of ear/tail DNA. The animals were housed in ventilated cages, maintained on a 12-hour light/dark cycle, and provided *ad libitum* access to food and water. All the mice were individually identified by ear tagging and their general health was closely monitored. All experiments involving mice were conducted with prior approval from the ethical committee at UCLouvain (2021/UCL/MD/018).

### Tissue harvesting

All the mice were sacrificed by cervical dislocation. The cranium was opened, and the brain was cut sagittally in half. Next, the two OBs and the two hippocampi were collected. The mouse snout was then recovered and cut along the midsagittal plane between the two nasal cavities. The OE was recovered from these cavities, distinguishable by its characteristic yellow color. The harvested tissues were snap-frozen by immersion in isopentane maintained at freezing temperature and then stored at –80°C until use.

## Sample preparation for immunostaining

### Decalcification

For immunostaining experiments, decalcification was needed to preserve the OE structure because of its localization within the nasal cavity. The skin, eyes, and lower jaw were removed from the mouse head. The head was directly rinsed in phosphate-buffered saline (PBS) and subsequently fixed with modified Davidson’s fixative (mDF) (37% formalin, ethanol, glacial acetic acid, water) for four days under agitation at room temperature (RT). The heads were rinsed for 24 hours in PBS and transferred to OsteoRAL R (RAL Diagnostics, #320715-2500), a commercial decalcifier, for five days under agitation at RT. After decalcification, the mouse heads were embedded in paraffin.

### Paraffin sections

Paraffin-embedded sections were obtained via a microtome. Coronal sections of 5 μm and 10 μm thickness were placed on Superfrost slides for IF and IHC, respectively.

### Immunohistochemistry, histological staining, and immunofluorescence

For immunohistochemistry, paraffin-embedded sections of 10 μm were immersed in different solutions as follows: 2×5 minutes in toluene, 2×2 minutes in absolute isopropanol, and 30 minutes in 0,3% H_2_O_2_ in methanol. The slides were finally placed in distilled water. Then, 1X Tris-buffered saline (TBS) was placed on each paraffin section for 10 minutes. This washing step was repeated two times.

Then, the sections were blocked with 10% normal goat serum (NGS) diluted in TBS for one hour at room temperature. Primary antibodies of interest were prepared in 1% NGS (supplementary table S1), placed on each section, and left overnight at RT. The negative control was incubated with 1% NGS only. The next day, the sections were rinsed three times for 10 minutes with TBS 1X. The secondary antibodies were diluted with TBS 1X and put on the corresponding sections for 30 minutes at room temperature. This step was followed by 3 TBS rinses. Stabilized Elite ABC reagent (Vector Laboratories, #PK-7100) was added to each section for 30 minutes. After another washing step, ImmPACT® DAB EqV Substrate (Vector Laboratories, #VEC.SK-4103-100) was prepared. Equivalent amounts of reagent 1 and reagent 2 were mixed and incubated on each section. The reaction was stopped after three minutes by transferring the slides into distilled water. The sections were dehydrated by immersion in different solutions as follows: 2 minutes in 70% isopropanol, 2 minutes in 90% isopropanol, 2×2 minutes in absolute isopropanol and 3×2 minutes in toluene. The sections were mounted with DPX mounting medium (Sigma-Aldrich, #1.00579.0500) and left to dry overnight at room temperature. The images were captured via an Axioskop 40 microscope (Zeiss).

Paraffin-embedded sections of 10 μm were stained with the Gallyas silver-staining method to identify NFTs (Alzheimer and Other Tauopathies Research Group, ULB Neuroscience, Karelle Leroy)[26]. The images were captured via an Axioskop 40 microscope (Zeiss).

For immunofluorescence, 5 μm paraffin sections were dewaxed by immersion in different solutions as described below: 3×5 minutes in toluene, 3 minutes in absolute isopropanol, 3 minutes in 100% ethanol, 3 minutes in 90% ethanol, 3 minutes in 70% ethanol, 2 minutes in distilled water and a final rinse in 1X PBS. Heat-induced antigen retrieval was performed using 10 mM sodium citrate buffer at pH 6.0. The sections were placed in a tank filled with antigen retrieval solution and heated for 3×5 minutes in a microwave, maintaining the temperature around 96°C. The solution was left to cool for 20 minutes, followed by rinsing with PBS.

For the permeabilization step, the sections were immersed in 0.3% Triton X-100 in PBS for 30 minutes at room temperature. The blocking solution was prepared with PBS-Triton, normal goat serum (NGS) and bovine serum albumin (BSA) to reach 0.3% PBS-Triton, 3% NGS and 5% BSA. This solution was added to each section for one hour at room temperature. The primary antibodies were prepared in blocking solution (Antibodies table S1), added to the corresponding sections, and left overnight at 4°C.

The next day, 3×5 minutes rinses with PBS-Triton 0.1% were performed, followed by one-hour incubation at room temperature with the secondary antibodies prepared in in 0.1% Triton X-100 in PBS (Antibodies table S1). The sections were rinsed twice in 0.1% Triton X-100 in PBS for 5 minutes, followed by a final rinse in 1X PBS. The sections were mounted with Mowiol (Merck, #475904-100GM-M) and left to dry overnight. The images were acquired via an EVOS^TM^ FL Auto fluorescence or confocal microscope.

All immunostainings were performed in PS19 males, as sexual dimorphism has been described in this model, with males exhibiting an earlier onset of the pathology [58].

### Behavioral tests

All behavioral tests were approved beforehand by the ethical committee (2021/UCL/MD/018) and conducted in collaboration with the behavioral analysis platform (BEAP, UCLouvain) following validated protocols. A dedicated cohort of mice (males and females) was used to perform olfactory function measurements at 3, 6 and 9 months. One week before the required time point, the mice were acclimatized on the behavioral analysis platform. Each mouse performed the olfactory discrimination test followed by the buried food test at each time point. Between the analyses, the mice were kept in quarantine and monitored closely. At 9 months after the last test, the mice were sacrificed by cervical dislocation, and the tissues of interest were harvested.

### Olfactory discrimination test

Test animals were placed in a clean plastic cage with a fresh layer of bedding covered by a removable Plexiglas lid with a 1.25 cm central hole for odorant exposure. After a five-minute adaptation period, a dry disposable applicator with a cotton tip was placed at a depth of 2.5 cm through the hole in the cage lid. After each minute, the applicator was removed and immediately replaced with a new applicator. After the sixth minute, an applicator with a cotton tip saturated with an odorant solution was positioned through the hole and again replaced every minute by a fresh applicator containing the same odor. After the 12^th^ minute, an applicator with a tip saturated with a different odorant was inserted and replaced six times at one-minute intervals (see timeline below). The two odors used were geraniol and lime. The number of times the animal headed toward each applicator introduced into the cage was recorded via a camera throughout the experiment and was counted by an experimenter blinded to the experimental conditions.

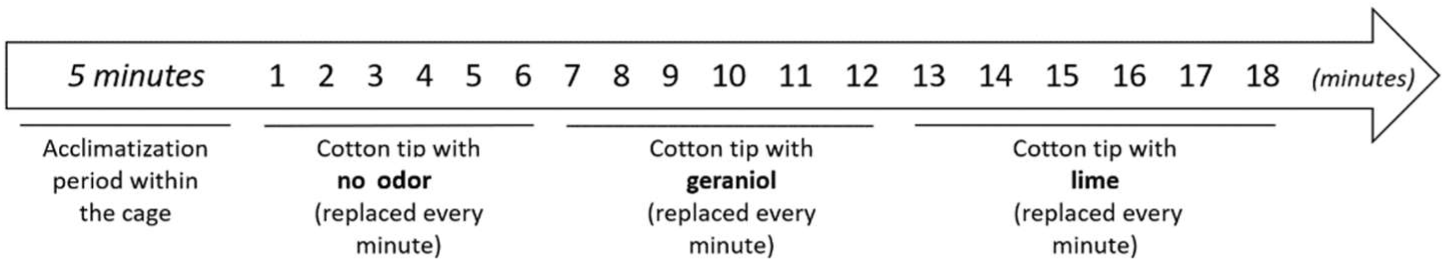

### Food-seeking test

The mice were deprived of food for 16 hours with access to water *ad libitum*. The test mice were introduced into a clean cage covered with a 3 cm thick layer of bedding, under which a 1.0-gram food pellet had been buried. The time elapsed between the animal’s introduction into the cage and the moment it retrieved the food with its front paws was measured in seconds, with a maximum of 300 seconds. From then on, a result equivalent to 300 seconds indicates that the mouse did not find the food.

### Intranasal ZnSO_4_ irrigation

At 1.5 months, the mice received an intranasal application of either 0.17M (5%) zinc sulfate (ZnSO_4_) or vehicle solution (PBS). Awake mice were held on their backs by one experimenter while a second experimenter administered 25 µL of either PBS or ZnSO_4_ solution, dropwise to the right nostril with a few seconds between each drop. After an hour to allow the mice to recover, the same procedure was repeated for the left nostril. The mice were then monitored for the next hour for any breathing discomfort. Since the OE regenerates over time [43], the mice were treated three times: at 1.5, 2.5 and 4 months. The mice were finally sacrificed at 6 months by cervical dislocation and samples were retrieved. For the evaluation of olfactory functions following ZnSO_4_ treatment, the treatment was performed only once at 3 months, and olfactory tests were conducted 24 hours after the treatment.

### NFT-like tau accumulations quantification

For each region of interest, the numbers of NFT-like tau accumulations visualized with anti-pTau immunostaining were counted in an equivalent area corresponding to the sum of 5 delimited areas placed in that same region (Fig. S11). The number of NFT-like tau accumulations per mm^2^ in the vehicle and ZnSO_4_-treated conditions was compared and analyzed via statistical tests.

### *Postmortem* human tissues

OBs were collected from 4 different Braak stage I or II patients and the OE was retrieved from 1 Braak stage II patient (Table S2). The autopsies were performed in UCLouvain (Belgium) in accordance with the applicable laws. Informed consent was obtained following local legislation. The recruitment protocols for the collection of human samples were approved by the ethical committees of UCLouvain (2020/02JUL/355). After collection, the tissues were immediately placed in 10% formalin for 48 hours. The tissues were subsequently embedded in paraffin and cut to obtain 5 and 10 μm paraffin sections. Neuropathological evaluation was conducted for each patient at KU Leuven (Leuven Brain Institute, Dietmar Thal), and the patients were classified based on the Braak stage (I or II in this study). Neuropathological analysis of these brains was approved by the UZ/KU-Leuven ethical committee (S-64363).

### Statistical analyses

GraphPad Prism software version 10 (GraphPad Software, Inc., La Jolla, CA, USA) was used for statistical analysis. All the data were tested for normality using the Shapiro–Wilk test. Parametric Student’s *t*-tests or nonparametric Mann-Whitney tests were used to evaluate the level of significance between two groups. When more than two groups were compared, parametric one-way or two-way ANOVA with the indicated post-hoc tests or nonparametric Kruskal–Wallis were used. The data are expressed as the mean +/− SEM. and P values <0.05 were considered significant (*P < 0.05, **P < 0.01, ***P < 0.001). The number of mice used in each group of experiment is indicated by “n=”.

## Results

### Tau pathology in the olfactory system and associated CNS regions in PS19 mice

The presence of hyperphosphorylated tau protein (pTau) was evidenced using the AT8 antibody (pTau (Ser202, Thr205)) in coronal sections of the OE from WT and PS19 mice. To that end, a specific protocol of decalcification has been set up to preserve the integrity of the OE. AT8 immunoreactivity was observed only in PS19 mice and was localized to the OE middle stratum and to the axon bundles (ABs), located in the lamina propria underlying the OE (Fig. 1a-d). The AT8 signal in the OE was assessed and compared at different ages: 1.5, 3, 6 and 9 months (Fig. 1a-d). AT8 immunoreactivity was detected as soon as 1.5 months (Fig.1a,a’), increased progressively up to 6 months (Fig. 1c,c’), and decreased at 9 months, especially in the middle stratum (Fig. 1d,d’). Based on morphology, no NFT-like tau accumulations were observed in this region for up to 9 months. A closer examination of the nasal cavity showed no pTau signal in the RE of PS19 mice (Fig. 1e). No pTau signal was detected in the OE or ABs of 6-month-old WT mice (Fig. 1f), indicating that the AT8 signal is specific to PS19 mice and related to the expression of pathological human tau. The results were confirmed independently by Western Blotting (Fig. S1) using the PHF-1 antibody. This antibody is generated against paired helical filaments isolated from the brains of AD patients, that recognizes tau phosphorylated at Ser396 and Ser404. It binds to tau filaments and is a recognized standard to evidence the pathological conformations found in NFTs [28, 48]. Immunostaining with the PHF-1 antibody and a human tau (hTau)-specific antibody showed similar results to those obtained with the AT8 antibody in PS19 mice (Fig. S2).

**Fig. 1.**
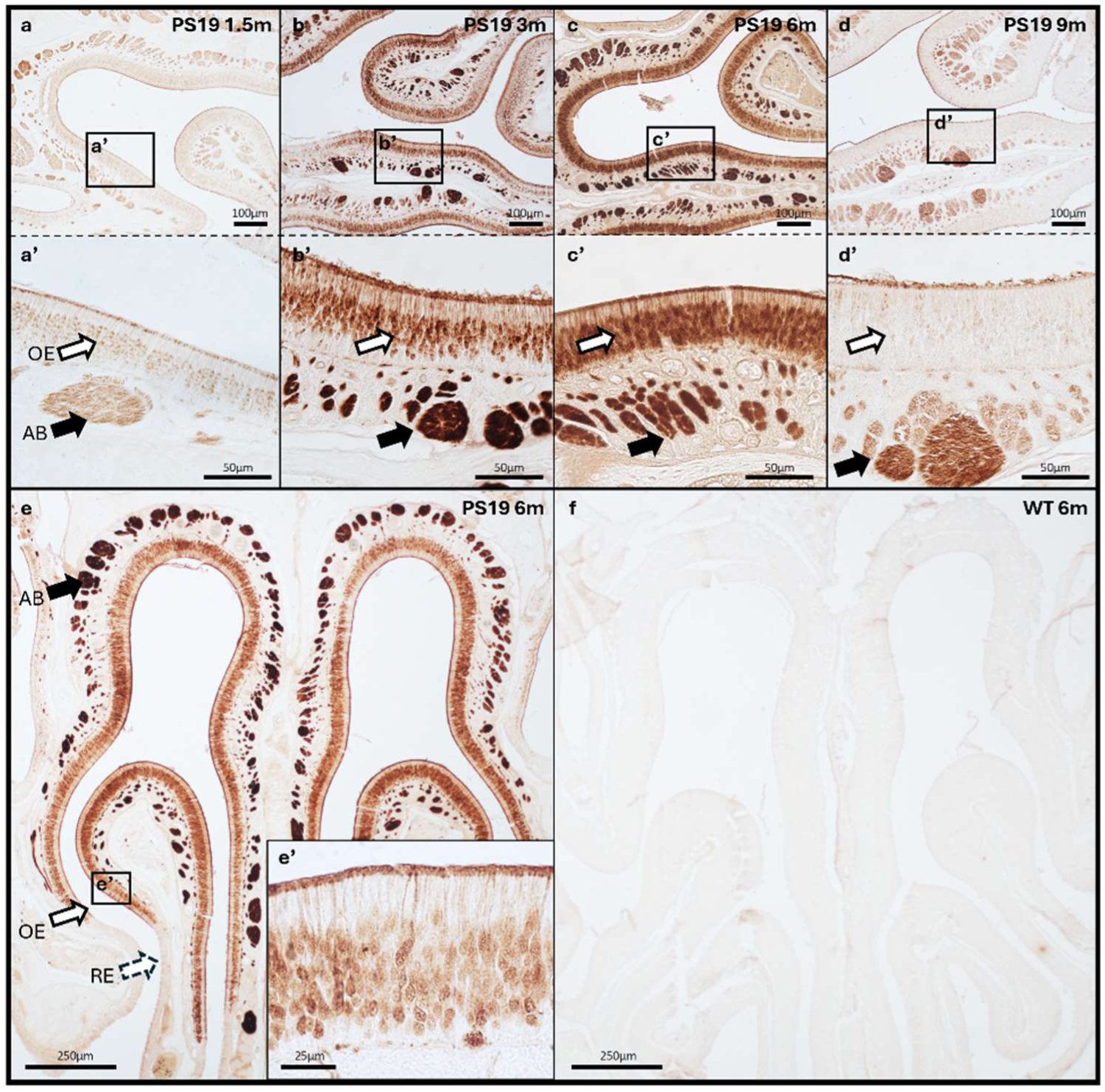
pTau expression in the OE of PS19 mice. The OE from 1.5-month (a,a’), 3-month (b,b’), 6-month (c,c’,e), 9-month-old PS19 mice (d,d’), and 6-month-old WT mice (f) were fixed and stained with the **AT8** antibody (pTau (Ser202,Thr205)). pTau recognized by the **AT8** antibody is found in the middle stratum of the OE (white arrows) and in the ABs (black arrows) as soon as 1.5 months in PS19 mice (a,a’). The signal increases up to 6 months (c,c’,e) and appears to decrease at 9 months (d,d’). No signal is detected in WT mice (f). All the mice used were **males**. The immunostainings shown are representative of n=4 mice per time point. High magnification of the OE (white arrows) is shown in the inset (e’). ABs: axon bundles, OE: olfactory epithelium, RE: respiratory epithelium.

Additional staining with a total tau antibody, detecting both endogenous murine tau protein and human tau protein, confirmed the presence of endogenous tau in the OE of WT mice (Fig. S2). RT-qPCR experiments indicated that the expression of endogenous tau transcripts was stable from 3 to 9 months in the OE of WT mice (Fig. S3). Since tau aggregation does not naturally occur in WT mice, overexpression of human tau with aggregation-prone mutations is required to induce tau pathology in animal models. Together, these observations indicate (i) that tau is endogenously expressed in the mouse OE, (ii) that the specific AT8-positive signal observed in the OE of PS19 mice is not an artifact simply related to ectopic expression of the human tau transgene in in OE of PS19 mice.

Colocalization analyses were carried out via immunofluorescence to confirm the cellular localization of the pTau signal in the OE of PS19 mice. The following antibodies were used: total tau antibody (TTau), olfactory marker protein (OMP) antibody (specific to mature OSNs and their neurites), and AT8 antibody (Fig. 2a-c). Colocalization was observed between the OMP signal and both TTau and AT8 signals in the cellular bodies and neurites of the OSNs, as well as in the ABs of the PS19 mice at 3 months (Fig. 2d,e). OMP and AT8 were also specifically colocalized in the dendritic knobs of OSNs (Fig. 2e). This result confirmed that pTau is found not only in mature OSNs of the OE but also in ABs within the lamina propria (Fig. 2e).

**Fig. 2.**
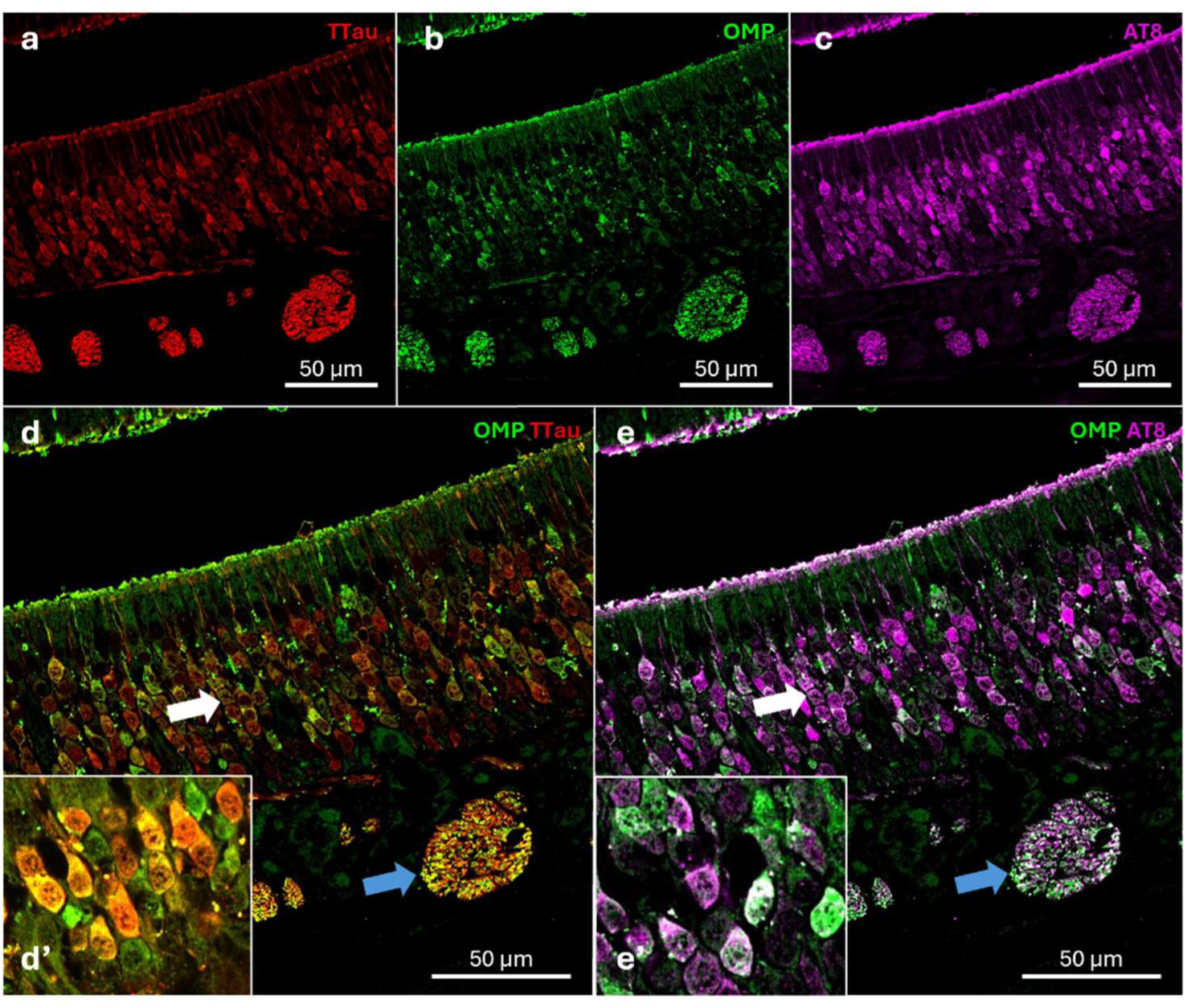
pTau expression in the OSNs and ABs of PS19 mice. Paraffin sections immunostained for total tau (**TTau**, red) (a,d), olfactory marker protein (**OMP,** green) (b,d,e) and pTau (**AT8,** purple) (c,e) show colocalization between **TTau-OMP** (yellow)(d) and **AT8-OMP** (white)(e) indicating tau hyperphosphorylation in the OSNs (white arrows) and in the ABs (blue arrows) of PS19 mice at 3 months. High magnifications (white arrows) are shown in the insets (d’,e’).

Since OSN axons extend through the cribriform plate of the ethmoid bone to reach the OB, where they form its outermost layer (ONL) [12], we next investigated the AT8 staining profile in the OB of PS19 mice. The AT8 signal was specifically present in the ONL of the OB as early as 1.5 months in PS19 mice (Fig. 3a). In addition, AT8 immunoreactivity was also detected in the MCL. The signal (ONL and MCL) remained stable for up to 6 months, indicating that pTau did not accumulate in these regions (Fig. 3a-d). In some 9-month-old mice, a stronger pTau signal appeared in the central region (e.g. GCL) of the OB (Fig. 3d). Here also, no AT8 signal was observed in the OB of WT mice (Fig. 3f).

**Fig. 3.**
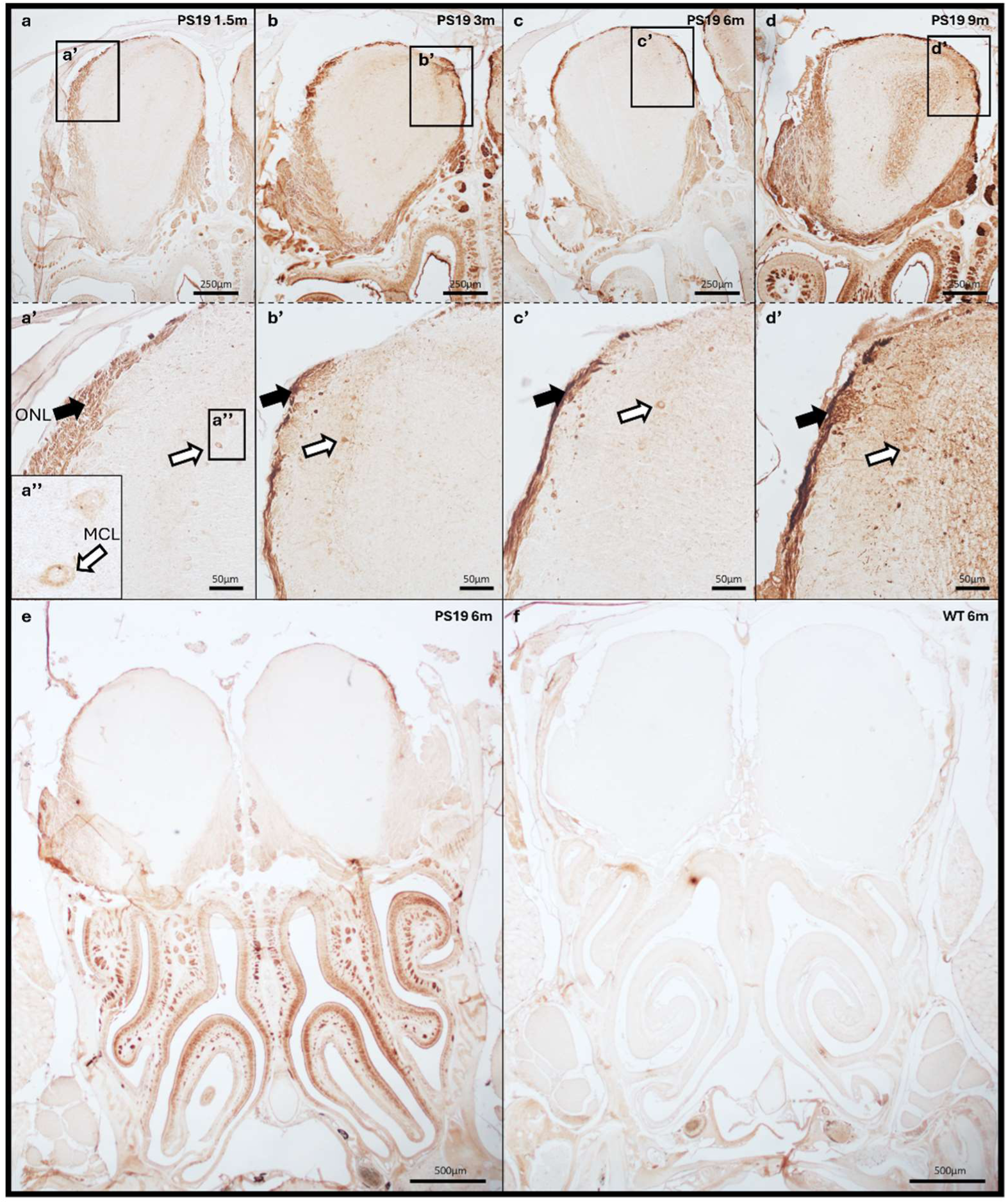
pTau expression in the OB of PS19 mice. The OB from 1.5-month (a,a’), 3-month (b,b’), 6-month (c,c’,e), 9-month-old PS19 mice (d,d’) and 6-month-old WT mice (f) were immunohistochemically stained with **AT8** (a-d, a’-d’). pTau recognized by the **AT8** antibody is found in the external layer of the OB (black arrows) and in the MCL (white arrows) from 1.5 months (a-d, a’-d’). The pTau signal seems stable over time (a’-d’). No signal is detected in WT mice (f). All the mice used for immunostainings were **males**. The immunostainings shown are representative of n=4 mice per time point. High magnification of the MCL (white arrows) is shown in the inset (a’’). ONL: olfactory nerve layer, MCL: mitral cells layer.

Accumulation of hyperphosphorylated tau in the OB of PS19 mice was confirmed by Western blotting of soluble protein extracts using the AT8 antibody (Fig. S4). The experiments were repeated and quantified using the PHF-1 antibody (Fig. S5). A band around 60kDa was observed exclusively in PS19 mice, both in males and females, as early as 3 months of age. No positive signal for PHF-1 or AT8 was observed in WT mice at any age (Fig. S5a-c). Western blot quantification revealed no significant differences in pTau expression between males and females in the OB at 3, 6 and 9 months (Fig. S5a-c). Finally, the PHF-1 signal was quite stable over time in PS19 males, suggesting that pTau, although already present at 3 months, did not further accumulate in the OB (Fig. S5d). The presence of human tau in the OBs of PS19 mice was confirmed by immunostaining with the Tau-13 antibody, as well as endogenous tau expression in the OBs of WT mice using a total tau antibody (Fig. S2). Endogenous tau expression in the OBs of WT mice was also confirmed by RT-qPCR (Fig. S6).

Immunofluorescence analyses of the OB in 3-month-old PS19 mice (Fig. 4a-c) revealed colocalization between TTau and OMP, indicating tau protein expression in both the ONL and the GL (Fig. 4d). Additionally, strong colocalization between OMP and AT8 was observed in the ONL, providing evidence of pTau accumulation in the outermost layer of the OB in PS19 mice (Fig. 4e). Some AT8 signal, which did not colocalize with OMP, was also observed in the inner layers of the OB (Fig. 4c).

**Fig. 4.**
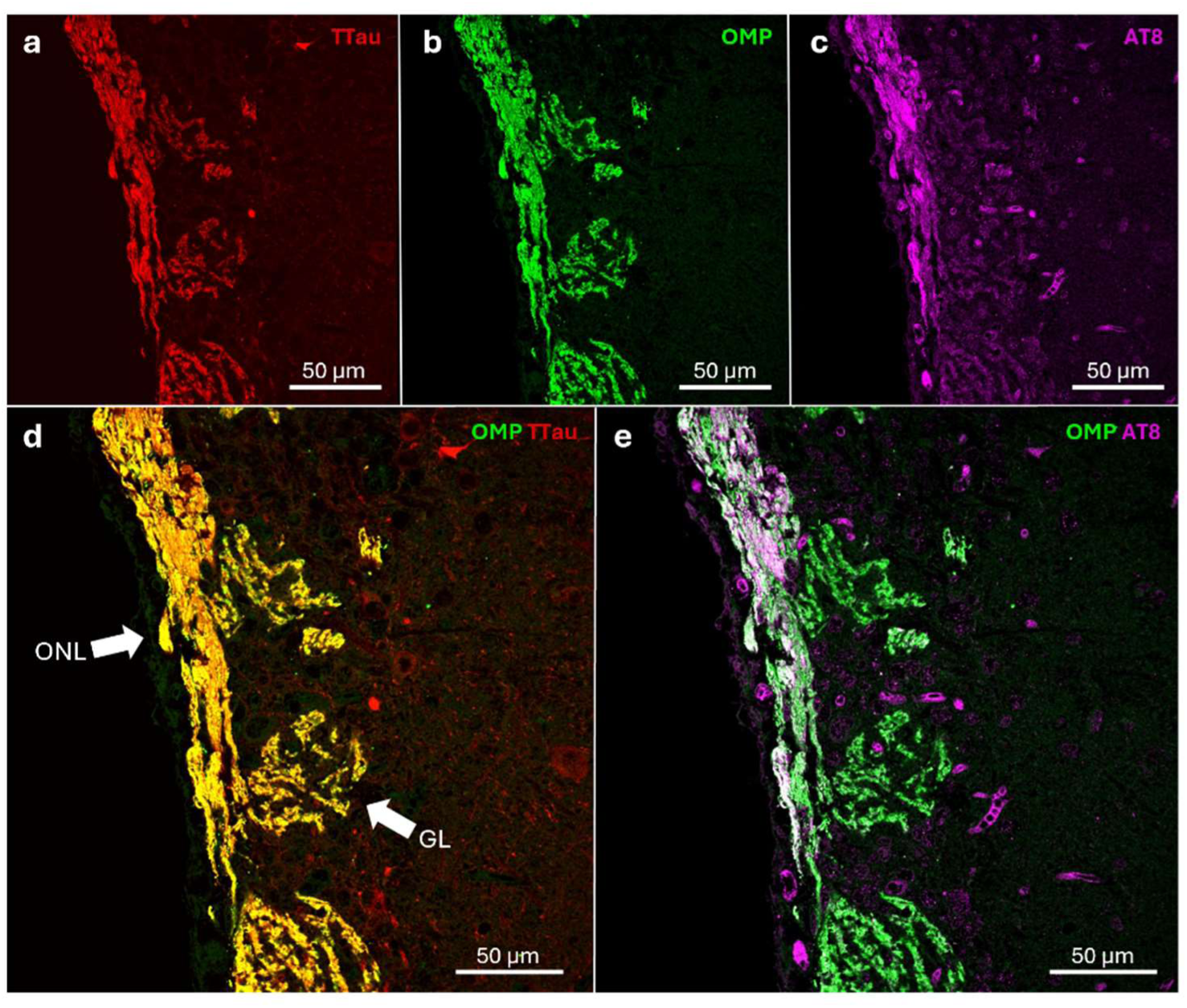
pTau expression in the ONL of PS19 mice. Paraffin sections immunostained for total tau (a,d) (**TTau**, red), **OMP** (b,d,e) (green) and **AT8** (c,e) (purple) show colocalization between **TTau-OMP** (yellow)(d) and **AT8-OMP** (white)(e) indicating tau protein expression both in ONL and GL as well as pTau expression in the ONL and inner layers of the OB from 3-month-old PS19 mice. ONL: olfactory nerve layer, GL: glomerular layer.

Next, we explored tau pathology in CNS regions of PS19 mice that are connected to the OB and involved in the processing of olfactory information along with olfactory memory: the PC, the EC, and the hippocampus. All these CNS regions are connected by neuroanatomical pathways [41]. Efferent projections from the OB reach both the PC and EC, and the perforant path wires the EC to hippocampal subregions [2]. In the PC, faint pTau immunoreactivity (AT8), indicative of the first step of abnormal tau modification [3], appeared in the neuronal cell bodies at 3 months (Fig. 5b,b’). By 6 months, the first perikaryal NFT-like tau accumulations were observed, with an accumulation of pTau in both the cell bodies and neurites of neurons within the PC pyramidal layer (Fig. 5c,c’). These tau accumulations progressively increased up to 9 months and eventually spread throughout the entire PC (Fig. 5d,d’). 9-month-old WT mice showed no AT8-positive accumulations in the PC (Fig. 5e). In the EC, AT8 immunoreactivity appeared in the neuronal cell bodies as early as 1.5 months and increased over time (Fig. 5f-i). Perikaryal NFT-like tau accumulations were detectable at 6 months, with signal present in both the cell bodies and neurites in the lateral part of the EC (Fig. 5h,h’). Tau lesions gradually accumulated up to 9 months and spread to the whole EC (Fig. 5i,i’). The accumulation pattern of tau pathology was similar to that observed in the PC, although with an earlier onset. In WT mice, AT8 signal was not detected until 9 months of age (Fig. 5j).

**Fig. 5.**
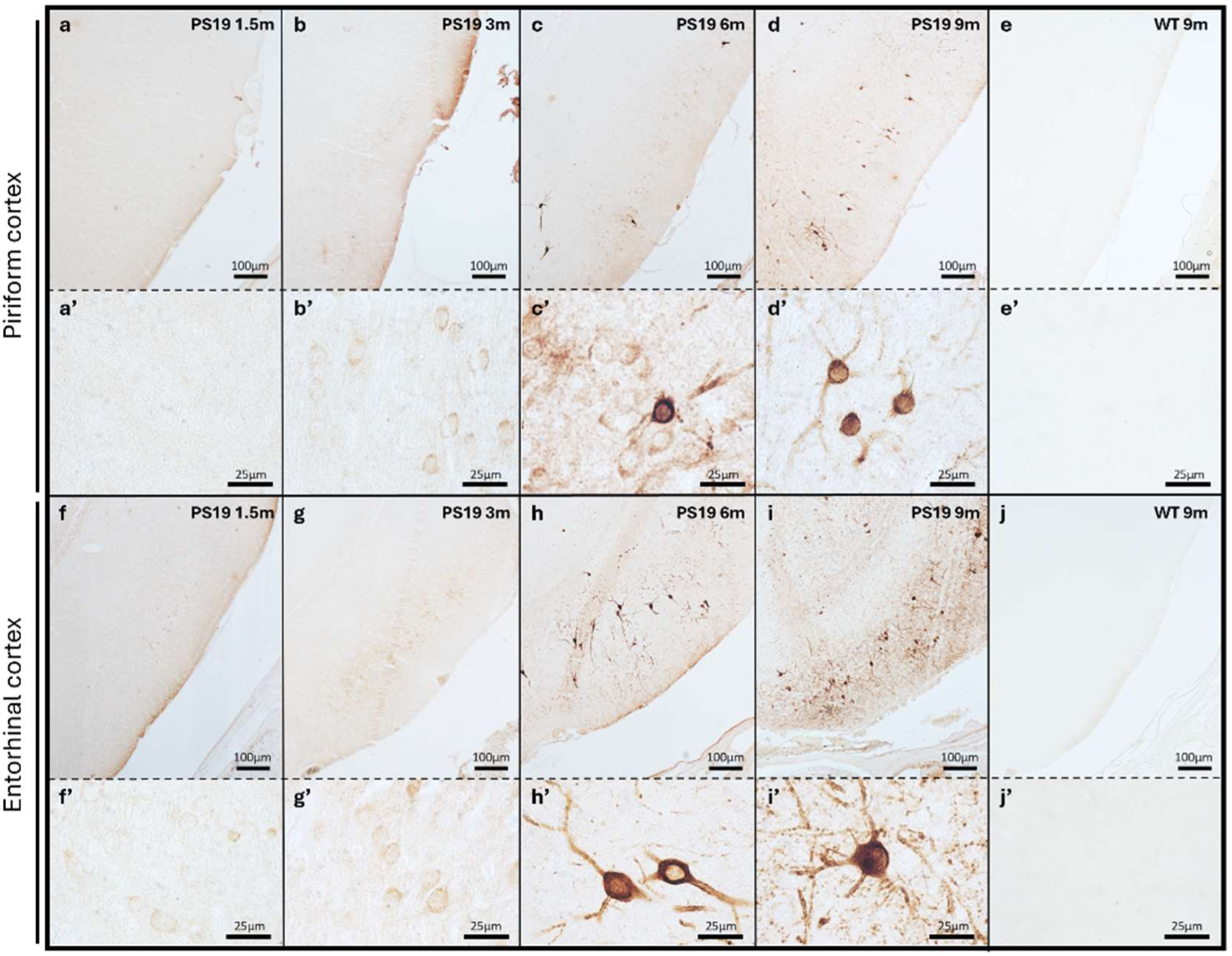
pTau expression in the PC and EC of PS19 mice. The PC from 1.5-month (a,a’), 3-month (b,b’), 6-month (c,c’), 9-month-old PS19 mice (d,d’) and 9-month-old WT mice (e,e’) were immunohistochemically stained with **AT8** (a-e). pTau recognized by the **AT8** antibody is found in the PC as soon as 3 months (b,b’) with an accumulation up to 9 months (b-d,b’-d’). The first NFT-like tau accumulations appear at 6 months (c,c’). There is no **AT8** signal in the PC of WT mice (e,e’). The EC from 1.5-month (f,f’), 3-month (g,g’), 6-month (h,h’), 9-month-old PS19 mice (i,i’) and 9-month-old WT mice (j,j’) were immunohistochemically stained with **AT8** (f-j). pTau recognized by the **AT8** antibody is found in the EC as soon as 1.5 months (f,f’) with an accumulation up to 9 months (f-i,f’-i’). PS19 mice develop NFT-like tau accumulations in this region at 6 months (h,h’). No **AT8** signal is detected in the EC of WT mice (j,j’). All the mice used were **males**. The immunostainings shown are representative of n=4 mice per time point. PC: piriform cortex, EC: entorhinal cortex.

In the hippocampus, AT8 immunoreactivity was faintly detected in the CA3 subregion at 3 months (Fig. 6b). By 6 months, pTau (AT8) was clearly present in the CA3 region and in the dentate gyrus (DG) (Fig. 6c), with an increased signal at 9 months (Fig. 6d). In the CA3 region, pTau was found in the CA3 mossy fibers and in the stratum pyramidale (Sp). In the DG, pTau was located primarily in the granular cell layer (Fig. 6e). No AT8 signal was detected in the hippocampus of WT mice (Fig. 6f).

**Fig. 6.**
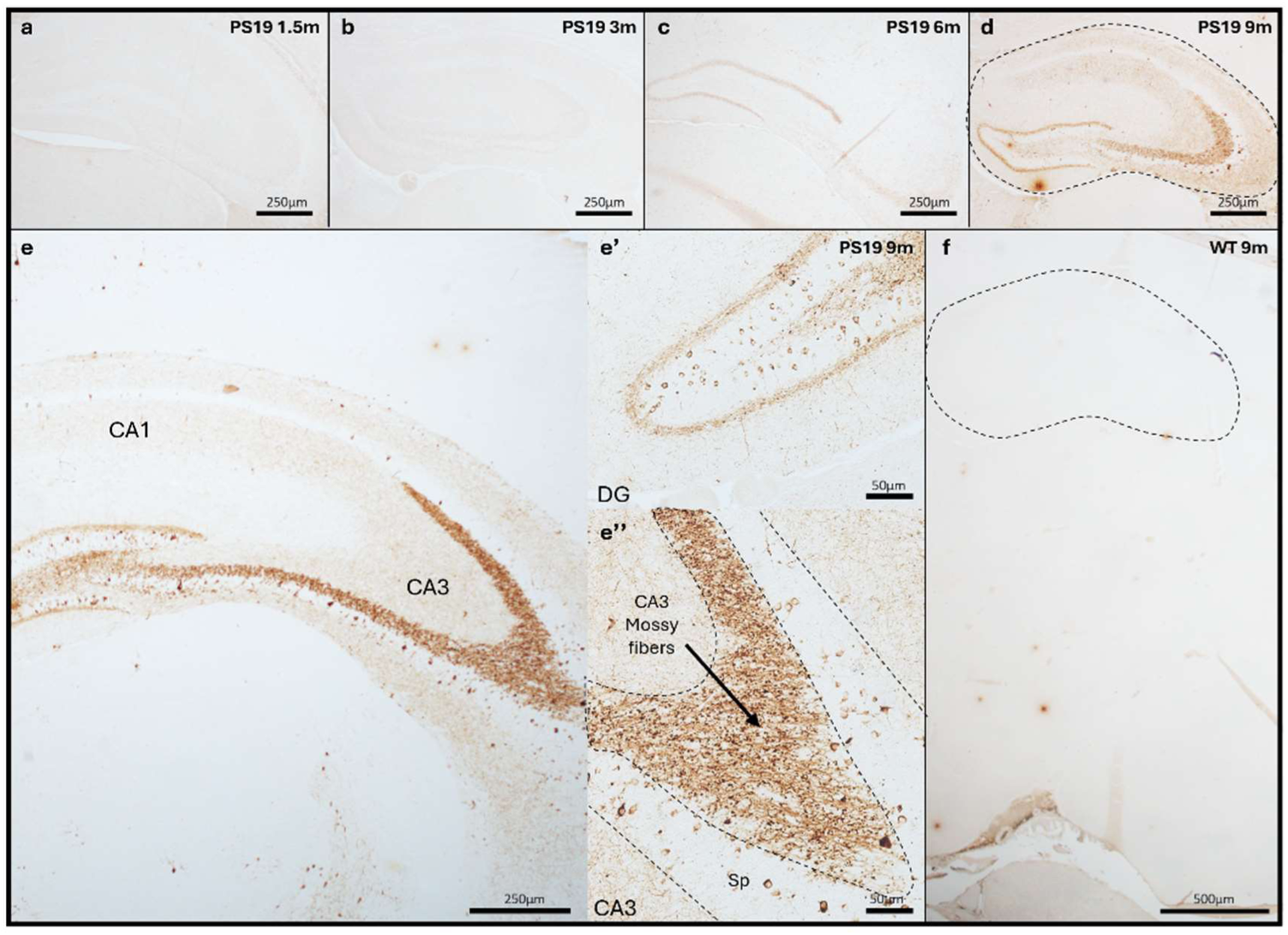
pTau expression in the hippocampus of PS19 mice. The hippocampus from 1.5-month (a), 3-month (b), 6-month (c), 9-month-old PS19 mice (d,e) and 9-month-old WT mice (f) were immunohistochemically stained with **AT8** (a-f). pTau recognized by the **AT8** antibody is found in specific hippocampal subregions like the CA3 (e’’) and DG (e’) as soon as 6 months with an increased signal at 9 months (c-e). No **AT8** signal is found in the hippocampus of WT mice (f). All the mice used for immunostainings were **males**. The immunostainings shown are representative of n=4 mice per time point. DG: dentate gyrus, CA: Cornu Ammonis, Sp: Stratum pyramidale.

pTau expression in soluble hippocampal extracts was confirmed by Western blotting with the AT8 antibody (Fig. S7). Equivalent results were obtained and quantified using the PHF-1 antibody. pTau was detected as a band around 60kDa and was present only in PS19 mice, as soon as 3 months of age in both males and females. No positive signal for PHF-1 or AT8 was observed in the WT mice regardless of the aging stage (Fig. S8a-c). At the same age, the average pTau levels were significantly greater in males than in females. Although the difference was already significant at 3 months, it was more pronounced at 9 months (Fig. S8a-c). Western blot analysis finally revealed a significant increase in the pTau signal between 3 and 9 months in PS19 males, consistent with the staining results, suggesting that pTau accumulation in the hippocampus occurs in a time-dependent manner (Fig. S8d).

To distinguish AT8-positive NFT-like tau accumulations from mature filamentous tau aggregates referred to as mature NFTs, Gallyas silver staining was performed in all previously studied regions and time points. No NFTs were observed in the OE (Fig. 7a-d) and OB (Fig.7 f-i) of PS19 mice. In the PC and EC, Gallyas-positive NFTs were detected only at 9 months of age in PS19 mice (Fig. 7n,s), and were very sparse compared to the AT8 signal observed previously (Fig. 5). In the hippocampus of PS19 mice, no NFTs were observed at any age (Fig. 7u-x). As a control, no NFTs were detected by silver staining in any region of interest in WT mice (Fig. 7e,j,o,t,y).

**Fig. 7.**
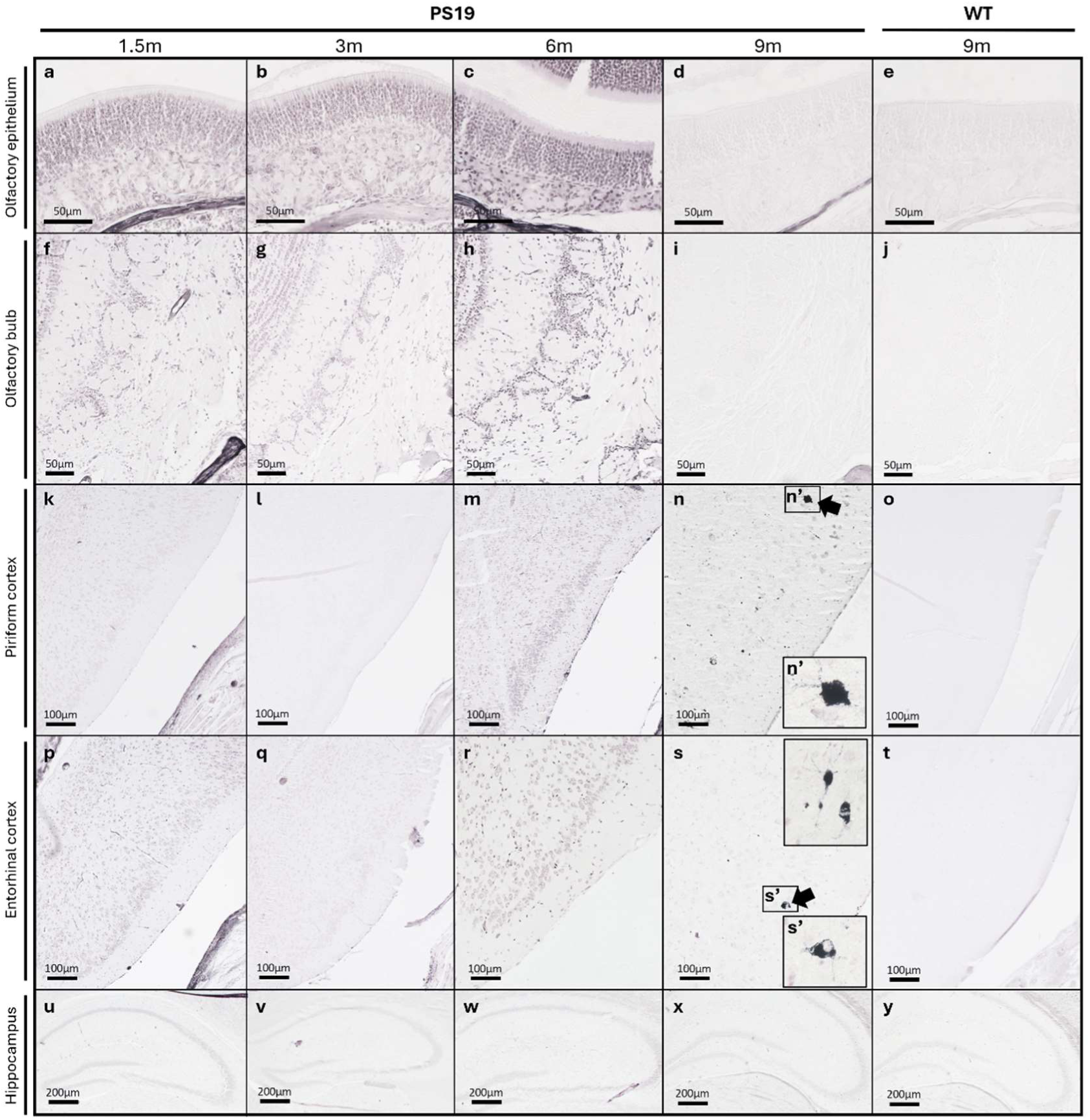
Gallyas staining in the OE, OB, PC, EC, and hippocampus of PS19 mice. The OE (a-e), OB (f,j), PC (k-o), EC (p-t) and hippocampus (u-y) of PS19 mice at 1.5, 3, 6, and 9 months, as well as 9-month-old WT mice were stained with Gallyas silver staining to highlight NFT inclusions. Few Gallyas-positive inclusions (black arrows) are only observed in the PC (n) and EC (s) from 9 months PS19 mice. No NFTs are detected in the OE, OB, or hippocampus at any time point, nor in the PC or EC before 9 months. No signal is detected in WT mice (e,j,o,t,y). High magnifications of the Gallyas-positive aggregates (black arrows) are shown in the inset (n’) and (s’). All the mice used for Gallyas silver staining were **males**.

These results clarify some aspects of the temporal and spatial staging of tau pathology in PS19 mice by distinguishing pTau (i.e. recognized by AT8 or PHF-1 antibodies), perikaryal NFT-like tau accumulations or pretangles (AT8-positive) and mature NFTs evidenced by Gallyas staining. Strikingly, pathological pTau accumulated very early (from 1.5 months of age) in the OE without the formation of typical NFTs, as shown by the Gallyas staining. A similar profile (pTau staining without NFTs) was observed in the OBs. pTau immunoreactivity was found in the EC and OE from 1.5 months on and in the hippocampus from 6 months. NFT-like tau accumulations appeared later, from 6 months in the PC and EC, and at 9 months in the hippocampus (Sp). While only a few NFTs (Gallyas-positive) were present in the PC and EC at 9 months. Taken together, our extensive analysis of the accumulation of pTau in connected olfactory regions gives a solid support to the hypothesis that tau pathology progresses along the olfactory system to reach regions of the temporal lobe that are particularly vulnerable in AD (e.g. EC, hippocampus).

### Olfactory function in PS19 mice

The impact of tau lesions in the olfactory system on olfactory function was assessed in PS19 mice using two specific olfactory tests: the olfactory discrimination test and the food-seeking test. The first test evaluated odor discrimination, whereas the food-seeking test measured odor threshold in mice. As a proof of concept, control and ZnSO_4_-treated mice were first evaluated since intranasal irrigation with ZnSO_4_ induces OE stripping and transient anosmia [44].

In the olfactory discrimination test, the curve for the number of sniffing events in non-treated (control mice) showed two peaks at 6 and 12 minutes (Fig. S9a). The first peak corresponded to the presentation of the geraniol odor, which induced a greater number of head movements toward the cotton swab in the control mice. Following this peak, the mice exhibited fewer head movements as they became accustomed to the odor. A second peak was observed after 12 minutes, when the second odor, lime, was presented. Both peaks were absent in the mice that received intranasal ZnSO_4_ administration (Fig.S9a). Similarly, during the food-seeking test, none of the ZnSO_4_-treated mice were able to locate buried food before the end of the test (300 seconds)(Fig. S9a’).

The same tests were conducted in WT and PS19 mice at 3, 6 and 9 months (Fig.8a-f) to evaluate the impact of tau pathology in the OE and associated CNS regions on olfactory function. In the olfactory discrimination test, both WT and PS19 mice showed two peaks corresponding to the introduction of each new odor. Each peak was followed by reduced reactions due to habituation to the odor (Fig.8a-f). All mice appeared to react more strongly, with a greater number of head movements, when presented with the second odor of lime. The superposition of curves indicates that all the mice, independently of their genotype and age, have similar olfactory capacities and that PS19 mice do not perform worse as they age. Statistical analyses revealed a significant impact of time as the mice responded to odor changes, and a not significant impact of genotype on olfactory capacities at 3, 6 and 9 months in both males and females (Fig. 8a-f). The results from the olfactory discrimination test suggested that the odor discrimination abilities of PS19 mice are not impaired by the presence of tau pathology in the OE. In the food-seeking test, there was no significant difference between WT and PS19 mice in the latency to find the pellet. This observation was consistent at 3, 6 and 9 months in both males and females. Compared to WT mice, PS19 mice did not exhibit an impaired ability to find the food (Fig. 8a’-f’).

**Fig. 8.**
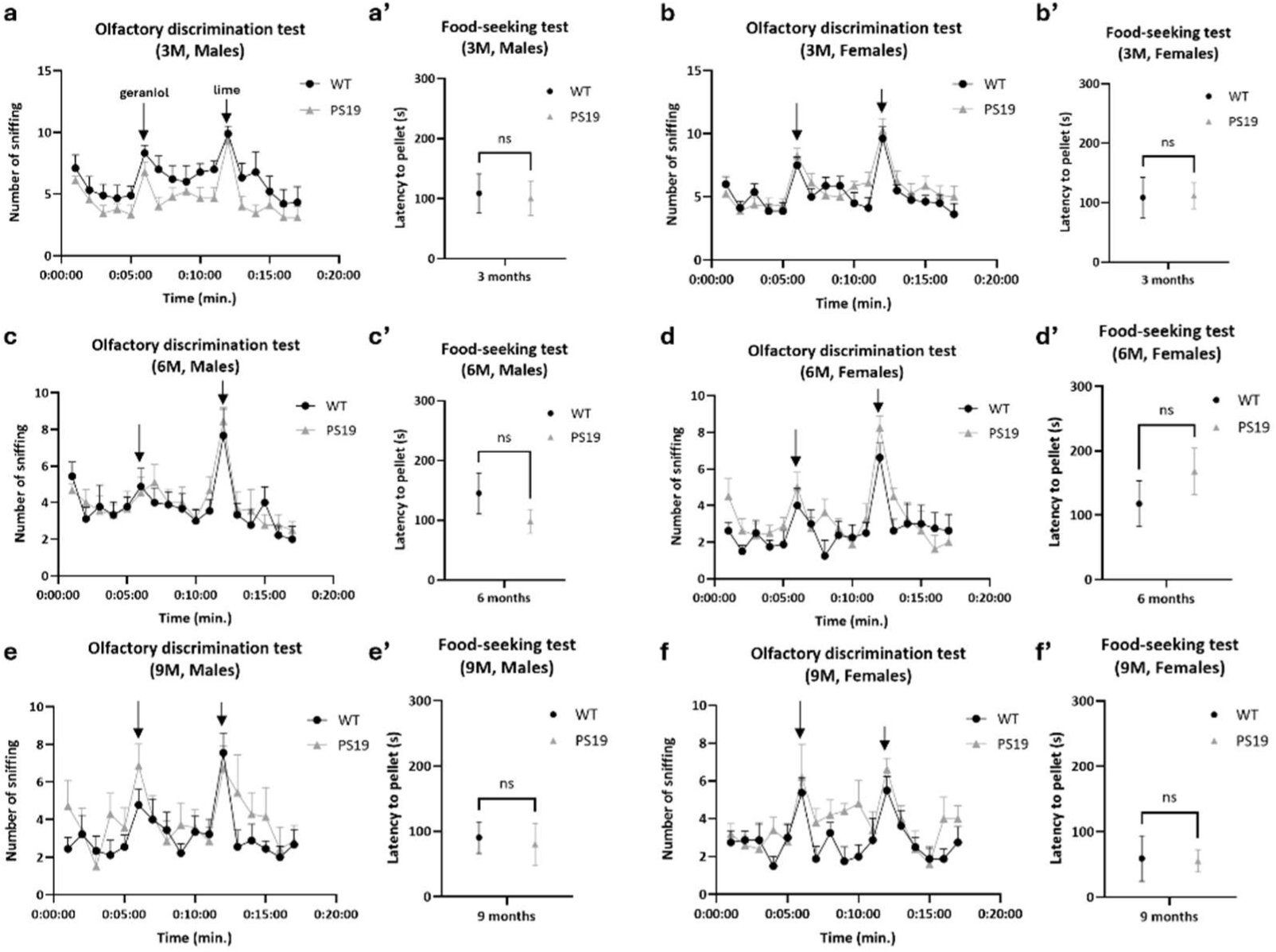
Olfactory function assessment in PS19 mice. The olfactory discrimination (a-f) and food-seeking tests (a’-f’) show no significant effect of genotype (PS19) on olfaction at either 3, 6 or 9 months, both in males and females (a-f, a’-f’). For PS19 males; n=9 (3 months), n=9 (6 months) and n=7 (9 months). For WT males; n=9 (3 months), n=9 (6 months) and n=9 (9 months). For PS19 females; n=8 (3 months), n=8 (6 months) and n=5 (9 months). For WT females; n=8 (3 months), n=8 (6 months) and n=8 (9 months). Data were analyzed using two-way ANOVA with Šídák’s multiple comparisons test (olfactory discrimination test) or using independent t-test (food-seeking test).

Together, these data indicate that despite progression of tau pathology along the olfactory regions with age, PS19 mice did not show a significant impairment in odor discrimination or detection threshold (sensitivity). Of note, another parameter (odor identification) difficult to address in mice was not evaluated here.

### Effect of intranasal ZnSO_4_ irrigation on tau pathology progression in PS19 mice

Given the apparent staging of tauopathy in olfactory regions and its early appearance in the OE, we examined the impact of targeted removal of the source of pathological tau present in the OE on tau pathology progression in the CNS. Our approach was designed to specifically remove the pTau accumulation in the OE using ZnSO₄ nasal irrigation, known to strip the OE and cause transient anosmia [43], which can later regenerate (OE) and recover (anosmia). Next, the impact on the progression of tauopathy in the OB, PC, EC, and amygdala (a connected region in the limbic system) was assessed. PS19 mice were intranasally administered either ZnSO₄ or phosphate-buffered saline (PBS, control). To ensure that the accumulation of tau in the OE was prevented for a sufficiently long period, nasal irrigation with ZnSO₄ was performed repeatedly at 1.5, 2.5, and 4 months prior to analysis of tauopathy at 6 months.

In vehicle-treated PS19 mice, the structure of the OE remained intact, with strong AT8-positive tau immunoreactivity in OSNs and ABs. However, following intranasal ZnSO₄ administration, the OE was completely stripped, and the AT8 signal was almost absent at 6 months (Fig. 9a). In the OB of PBS-treated mice, all six layers were present and structurally preserved, with AT8-positive signal characteristic of PS19 mice. After ZnSO₄ treatment, the AT8 signal in the ONL was no longer detectable (Fig. 9a). The efficient stripping of the OE and the ONL after ZnSO₄ treatment was further confirmed by Masson’s trichrome blue staining (Fig. S10a-d). Tau pathology was next examined in regions receiving direct inputs from the OB, such as the PC, EC, and amygdala. At 6 months, vehicle-treated PS19 mice exhibited NFT-like tau accumulations in these regions. However, following ZnSO₄ treatment of PS19 mice, no NFT-like tau accumulations were detectable in the PC, and only a few remained in the EC and amygdala at 6 months compared to control PS19 mice (Fig. 9a). Quantification of the number of NFT-like tau accumulations per mm^2^ in the PC, EC, and amygdala (Fig. S11), further confirmed a massive reduction of pTau signal associated to tau pathology in the PC, EC and amygdala following intranasal ZnSO_4_ administration (Fig. 9b). No overt morphological alterations were found in these regions after staining with Masson’s trichrome blue (Fig.S10e-j) indicating that ZnSO₄ treatment of the OE did not result in loss of density of the connected olfactory regions. In other terms, removing the source of pTau present in the OE sharply reduces tau pathology in the connected regions of the temporal lobe and limbic system. This indicates that early accumulation of pTau in the OE can be instrumental to the progression of tau pathology that eventually reaches the EC and the hippocampal formation.

**Fig. 9.**
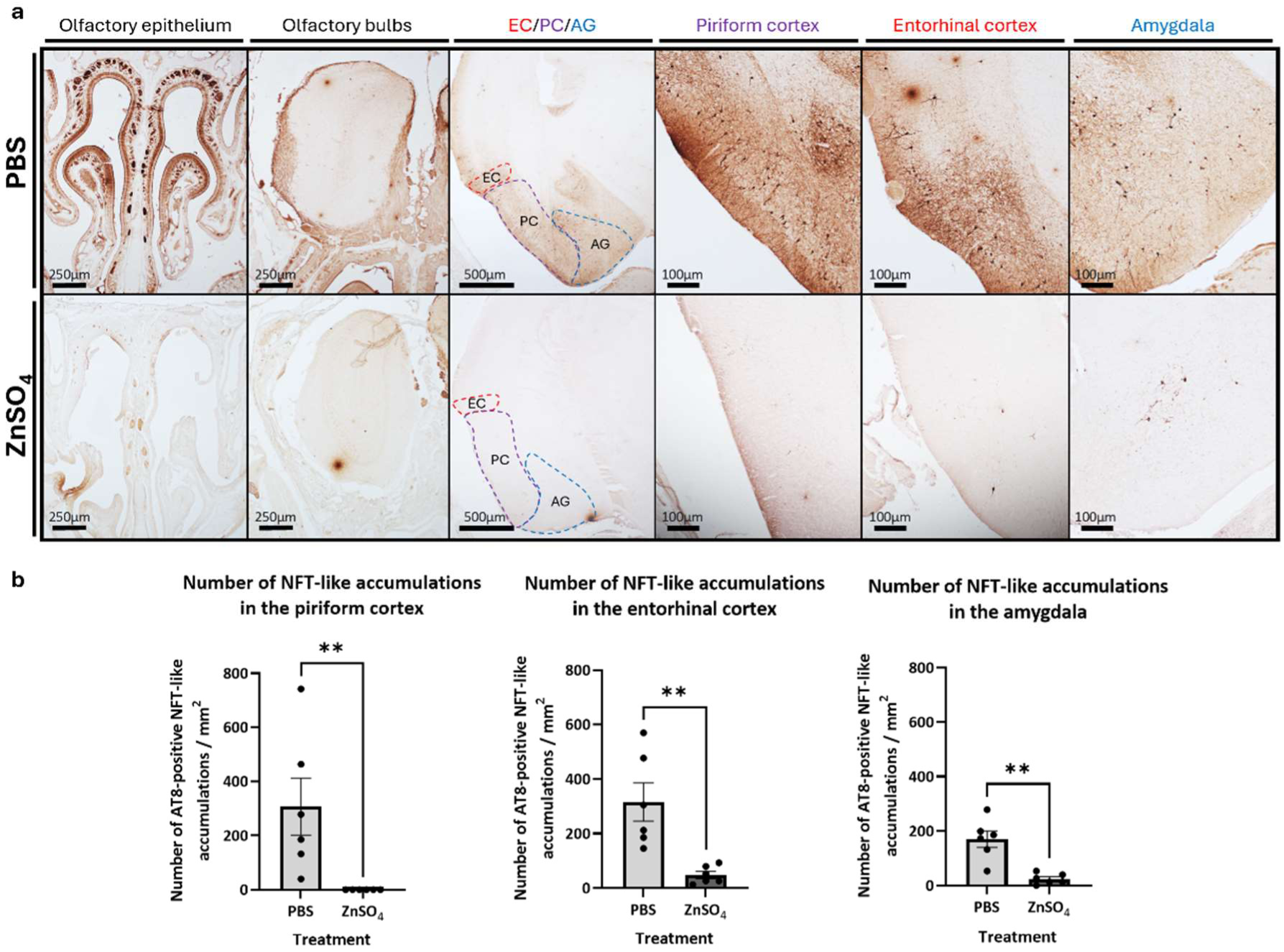
pTau expression in 6-month-old PS19 males after PBS or ZnSO4 nasal irrigation. Sections of the OE, OB, PC, EC, and amygdala from PS19 males were fixed and stained with the **AT8** antibody (a). After intranasal ZnSO_4_ irrigation, the OE and ONL from the OB are stripped, no more pTau signal is found in these regions. Quantification of the number of NFT-like tau accumulations/mm² was performed in the PC, EC, and amygdala (b). In the PC, there is no NFT-like tau accumulations compared to the PBS-treated mice. In the EC and amygdala there is a significant decrease in NFT-like accumulations compared to the control condition. **P < 0.01 (Mann Whitney test, n=6 mice/group).

### Pathological pTau protein in the human olfactory system

Finally, we investigated whether a similar pattern of tau pathology was observed in OE and OB collected from patients at early Braak stages I and II (patients 1-4, Table S2). In the OE collected from patient 4 (Braak stage II), pTau immunoreactivity (PHF-1 Ser396, Ser404) was detected and colocalizing with the OMP signal, indicating the presence of pTau in OSNs (Fig. 10b). PHF-1 signal was also detected in the ABs present within the OE structure and revealed by the OMP antibody (Fig. 10c-e). Interestingly, while colocalization was observed between OMP and PHF-1 within the ABs (Fig. 10f), AT8 signal was detected neither in the ABs (Fig. 10g) nor in the OSNs.

**Fig. 10.**
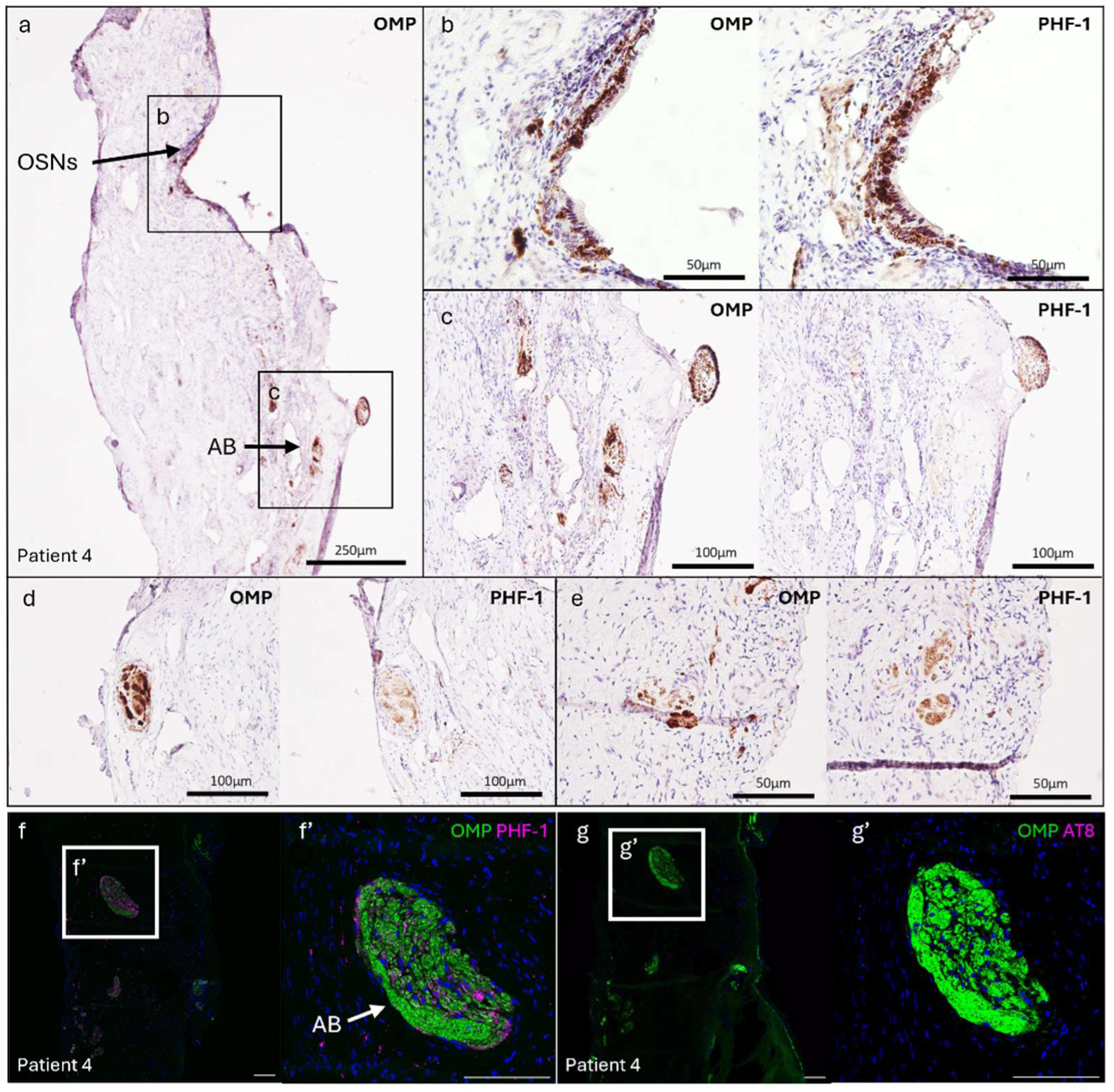
pTau expression in human OE. The OE was obtained from a Braak stage II patient (patient 4). The sections were immunohistochemically stained with **PHF-1** or with **OMP** (a-e). **PHF-1** signal is observed in the **OMP**-positive OSNs (b) and ABs (c-e) of the OE. Paraffin sections immunostained for **OMP** (green) and **PHF-1** (purple) show colocalization between **OMP-PHF-1** (white) in the AB (f,f’). PHF-1 signal is also observed inside the tissue (f). Paraffin sections immunostained for **OMP** (green) and **AT8** (purple) show no colocalization between **OMP-AT8** (g,g’). No signal is detected in the negative controls without primary antibodies. Scale bar: 100μm.

In the OB of Braak stage I/II patients, PHF-1 and OMP colocalization was consistently detected in the ONL, and to a lesser extent, in the GL (Fig. 11a-d,d’). Some PHF-1 pTau signal was additionally found in the inner layers of the OB (Fig. 11a-d,d”). No colocalization was observed between OMP and AT8 in the ONL (Fig. 11e-h,h’) but similarly to PHF-1, neuropil threads highlighted by the AT8 antibody were observed in the inner layers of the OB from Braak stage I and II patients (Fig. 11e-h,h”). These results underscore the early appearance of pathological tau markers in the sensory regions of the human olfactory system in agreement with observations in the PS19 mouse model.

**Fig. 11.**
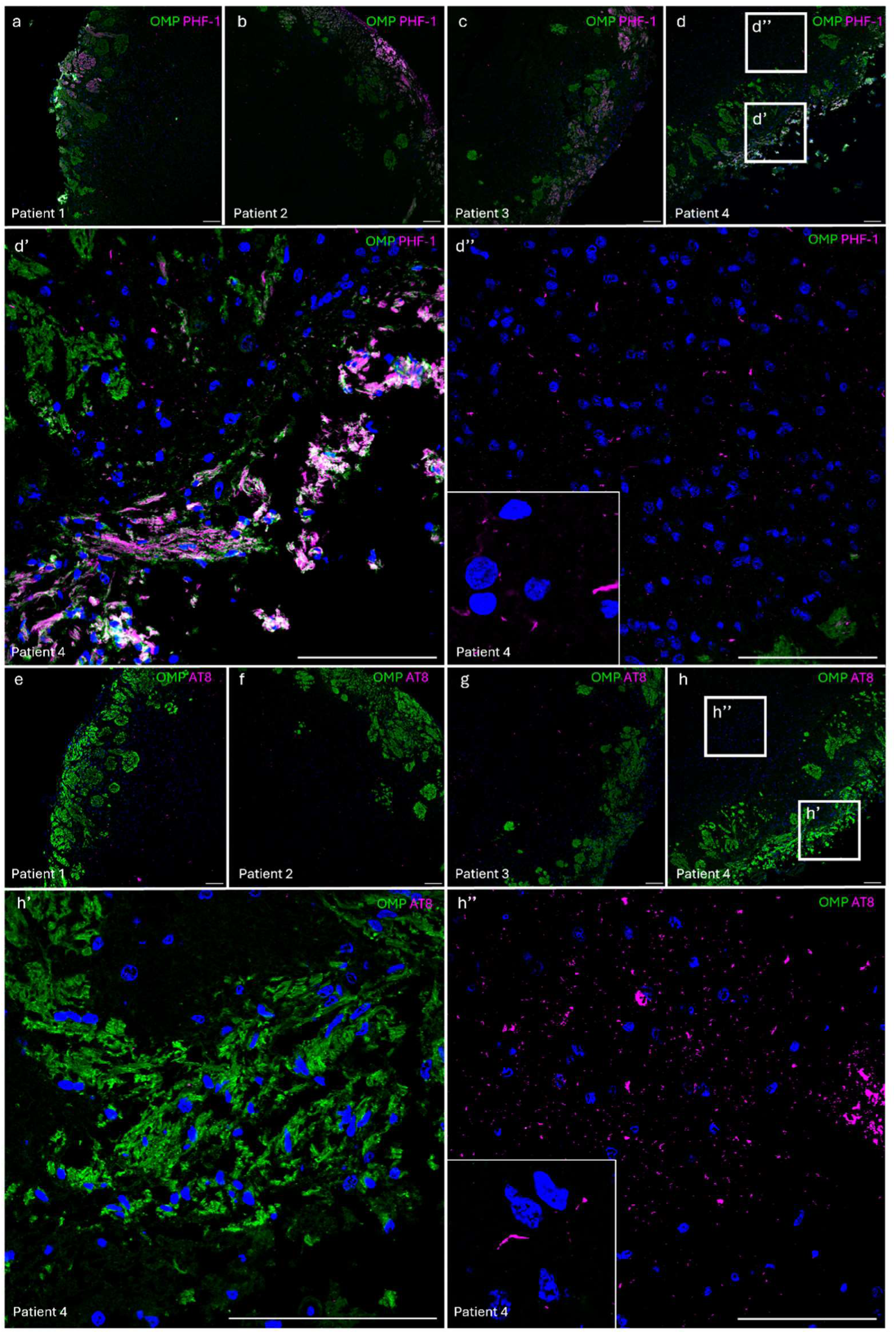
pTau expression in human OB. The OB were obtained either from patients 1 and 2 (Braak stage I)(a,b,e,f) or from patients 3 and 4 (Braak stage II)(c,d,g,h). Paraffin sections immunostained for **OMP** (green) and **PHF-1** (purple) show colocalization between **OMP-PHF-1** (white) in the ONL of all patients (a-d,d’). **PHF-1** signal is also found in the inner layers of the OB (a-d,d’’). Paraffin sections immunostained for **OMP** (green) and **AT8** (purple) show no colocalization between **OMP-AT8** (e-h,h’). AT8 signal is only found in the inner layers of the OB, similarly to **PHF-1** (e-h,h’’). No signal is detected in the negative controls without primary antibodies. Scale bar: 100μm.

## Discussion

The OE forms a direct appendix of the CNS. In fact, OSNs are the only neurons connected to the CNS being in direct contact with the external environment [57], lacking the protection of the blood-brain barrier [9]. These unique anatomical characteristics make the OE highly vulnerable to harming environmental conditions and a possible starting point for the spread of pathologies through its connections to various regions of the olfactory and central nervous systems. It also sheds light on the importance of studying this region from both diagnostic and therapeutic perspectives. Regarding tauopathies, the presence of pTau immunoreactivity in the OE of AD patients has been reported [4, 55, 57, 61]. NFTs are also present in the OB of AD patient early in AD progression [35, 46] and are correlated to bulb size reduction [42]. Tau pathology could thus appear early in the sensory part of the olfactory system (OE), due to neurotoxic compounds present in the environment or to the gradual loss of its ability to regenerate. Considering that tau spreading along neuronal connections is now well documented [51], one can imagine that pathological forms cross the cribriform plate to penetrate the CNS and spread through the olfactory tract to regions known to be affected in AD (EC, amygdala, hippocampus). Here, we provide the first evidence of the staging of tau pathology across the olfactory system, and, more importantly, that preventing the accumulation of pathological tau in the OE reduces NFT-like tau accumulations in the PC, EC, and amygdala.

### Prion-like spreading of tauopathy within the olfactory system

Our study revealed the presence of hyperphosphorylated tau protein in the OSNs of the OE as early as 1.5 months in PS19 mice. The signal increased up to 6 months, and decreased at 9 months, which could be explained by the reduction in the number of OSNs with aging [22]. Neurogenesis occurs in the OE throughout life but aging and exposure to pathogens contribute to a reduction in this regenerative capacity [14, 49]. pTau immunoreactivity was also detected in ABs suggesting that the signal appears early in OSNs and spreads following their axonal projections to accumulate within ABs. These nerve fibers then reach the OB following efferent projections and constitute its most external layer, the ONL. Immunohistochemical analyses showed strong pTau signal in the ONL but also in the MCL. Mitral cells in the MCL are neurons that receive afferences from OSNs within the olfactory glomeruli and project to various regions in the brain, notably the PC or EC that are part of the olfactory cortex. The OC allows the integration of olfactory information [9, 56]. pTau was also detected in these two cortical regions: as early as 3 months in the PC and 1.5 months in the lateral EC. Finally, the last region investigated was the hippocampus due to its connections with the EC layers II and III, which project to specific hippocampal subregions. Neurons from the entorhinal layer II project to the CA3 and the dentate gyrus while neurons from the entorhinal layer III project to the CA1 [64]. pTau immunoreactivity was observed in these two hippocampal subregions. The presence of tau pathology in PS19 mice was demonstrated in all areas of interest, from the OE to the hippocampus, and its time-dependent spread seemed to follow neuroanatomical pathways consistent with previous findings in the CNS regions of PS19 mice [51]. It was suggested that tau pathology may spread along neuroanatomically connected pathways. For instance, injection of brain extracts from PS19 mice into the brains of transgenic animals expressing human tau animals induced the assembly of WT human tau into filaments and the spread of pathology from the injection site to neighboring brain regions [13]. This type of neuron-to-neuron progression is found in many cerebral proteionopathies, which share similar cellular and molecular patterns of pathological protein propagation [21]. Termed “prion-like” propagation, this process implies that the tau protein can actively propagate trans-synaptically and that misfolded tau can affect tau folding in healthy surrounding neurons through a seeding mechanism [21]. Owing to its direct connection with the OE and other CNS regions, the OB is often regarded as a hub for the spread of misfolded proteins or as an entry point for pathogens [57, 68, 70]. Our results suggest that pTau appears early in OSNs from the OE of PS19 mice and likely spreads following the olfactory pathway to the OB, without maturation of tau seeds into mature NFTs in these regions, as shown by the Gallyas staining. pTau then appears in the olfactory cortex (PC and EC) and further in the hippocampal regions (essentially CA3 and DG) which are projection regions of neurons from layer II (and possibly layer VI) of the EC. Of note, Gallyas-positive inclusions are readily present only in PC and EC 9 months, indicating that NFTs are a late manifestation of tau pathology that do not appear in all brain regions where pathological tau (e.g. AT8-positive) is available. This crucial point should be further investigated to define what pathological forms of tau are responsible for its transsynaptic spreading. Evidence from studies in the retina suggests that the induction of fibrillar aggregates is essential to induce pTau spreading in the brain [63]. Additionally, it is imperative to elucidate how pTau contributes to the formation of typical paired helicoidal filaments (PHFs) in more specific subregions.

Given the anatomical proximity between the OB and limbic regions, and considering the olfactory deficits in AD patients, it has previously been suggested that tau pathology spreads from olfactory circuits to limbic structures including the amygdala, as well as the piriform and entorhinal cortices [1, 54]. This route of propagation may exacerbate an already existing tauopathy within the CNS, making the OE a promising target region for early treatment intervention. The expression pattern of tau pathology observed in the olfactory system of PS19 mice might resemble that found in AD patients where tau-immunoreactive fibers have been reported in the ONL [34], and tau-containing neurites were present in the OE [4, 34]. We showed here pTau staining (PHF-1) in the ONL – formed by the axons of olfactory neurons –from Braak stage I/II patients, as well as in OSNs and ABs within the OE. PHF-1 has been reported to detect phosphorylated tau in axons and the neuropil, preceding pretangle formation, while AT8 stains pretangles and is considered an early marker of tau pathology [3]. In our observations, PHF-1 staining was detected in the OE and in the ONL of the OB but also evidenced neuropil threads in the inner layers of the OB, whereas AT8 staining was restricted to these inner layers of the bulbs. This suggests that hyperphosphorylated tau may be present in the OE at early Braak stages without converting over time to aggregate, possibly due regenerative capacity of the OE that continuously produces newborn OSNs. Still, these pTau forms detected by PHF-1 in the OSNs could be heading for the ONL, which is formed by the axons of the OSNs. Consistent with our data, hyperphosphorylated tau then accumulates within the inner layers of the OB to form pretangles evidenced by both AT8 and PHF-1 staining. These results and the underlying scenario of pTau progression are consistent with other studies showing phosphorylated tau evidenced by AT8 or PHF-1 staining in the OB of AD patients [24, 55].

This strongly suggests that tau pathology may develop early in these regions. These findings extend beyond the transgenic mouse model, indicating the relevance of our observations to human pathology. While there is an ongoing debate regarding whether tau pathology in the olfactory system arises earlier, simultaneously, or subsequently than in the EC, our results suggest that the olfactory system may be a site of early tau pathology development in NDs [34, 62]. Importantly, the direction of the progression of tau pathology in the olfactory system is matter of different hypothesis. On one hand, different studies postulate that the olfactory system serves as an entry point for pathogens or as an access point for environmental aggressions, which can trigger pathological changes that would then spread throughout the brain via anterograde olfactory pathways [14, 20, 52, 57] Our data illustrate this hypothesis but are not at odds with the one supporting Tau pathology appears early in the brain, particularly in the EC, and then spreads to the olfactory system using the retrograde olfactory circuit [17].

### Olfactory dysfunction: a prodromal sign of dementia

Olfactory deficits have been observed and associated with NDs, including AD, since 1974 [17, 65]. Olfactory dysfunction is present not only in the early stages of AD but also in MCI patients [57] and is now recognized as a prodromal sign of dementia [9] and as a tool to predict MCI-to-AD conversion [36].

Since then, behavioral studies aimed at evaluating olfactory function have been conducted to characterize mouse models of these pathologies. In our study, the results of the food-seeking and olfactory discrimination tests did not show any significant difference between the PS19 and age-matched control mice. PS19 mice did not take longer to find the buried food and reacted in the same way as the control mice did when presented with two different odors, contrary to what had been previously demonstrated [39]. These results could be explained by the fact that the term “olfactory deficits/dysfunction/impairment” encompasses several functions, such as a decline in odor threshold, odor identification, odor discrimination, and odor memory that cannot be measured by the same tests [9]. In addition, differentiating olfactory deficits caused by physiological aging from those associated with NDs presents a challenge [22]. During normal aging, the OE, which contains OSNs, is gradually replaced by respiratory epithelium. This is accompanied by ossification and closure of the foramina in the cribriform plate. Together, these changes can disrupt the transduction of olfactory information from the OE to the OB and the olfactory cortex [9, 57] and lead to an increase in odor threshold, which relies solely on the integrity of the OE and OSN transduction [9, 33]. In contrast, olfactory memory relies on hippocampal integrity, whereas odor identification/discrimination relies on the integrity of central olfactory structures [22, 34]. In AD patients, it has been shown that odor identification is particularly impaired [9, 36, 57, 65]. This impairment reflects a cognitive deficit rather than a direct impairment of olfactory perception [23, 34, 36], since higher cognitive skills, such as verbal and semantic skills, are required [22]. Some odor identification tests have already been proposed, in which certain odors can differentiate AD patients from individuals with normal aging [57]. Our findings show that following intranasal ZnSO_4_ irrigation, which disrupts the peripheral olfactory system [31], mice are no longer able to find buried food, suggesting an increased odor threshold. Additionally, they are no longer able to differentiate between odors, indicating that their discrimination abilities could be impaired. The effectiveness of the tests conducted cannot, therefore, be questioned. These tests primarily evaluate peripheral damage at the level of the OE (odor threshold). It is thus possible that PS19 mice do not present strong peripheral impairment despite the presence of tau pathology in the OE. They might retain their ability to smell odors because of the lack of fibrillar tau (NFTs) in this region. However, the presence of lesions in the primary olfactory cortex (OB, PC, EC) might compromise olfactory discrimination and identification [9], two functions that are truly challenging to evaluate in mice. Consistent with our hypothesis, in Tg2576 mice – an amyloid mouse model – odor memory is impaired while odor perception is spared, suggesting that the underlying mechanisms might be different [9]. In addition, it is known that, compared with mice, humans have a poorly developed sense of smell [38]. In rodents, the OE covers 50% of the nasal cavities, while it covers only 3 to 5% of these cavities in humans. Consequently, olfactory neuronal loss has a greater impact on odor perception in humans than in rodents [19]. Mice rely heavily on their olfactory function for survival, and given their significant olfactory capabilities, we hypothesize that the pTau pathological pattern detected in the olfactory regions might not be sufficient to induce olfactory dysfunction detectable by the two tests conducted.

### Targeting the OE mitigates tau pathology in the CNS

To establish a causal link demonstrating that the OE serves as a gateway for pathology and worsens pre-existing conditions in the CNS of PS19 mice, we directly targeted the OE and assessed its impact on the tau pathology progression in the CNS. Intranasal ZnSO_4_ irrigation was used to specifically strip the OSNs in the OE and ONL in the OB [31, 37], providing a model to study the downstream effects on the CNS. Repeated intranasal irrigation with ZnSO_4_ in PS19 mice resulted in near-complete stripping of the OE and ONL. Additionally, NFT-like tau accumulations were absent in the PC and only a few residual lesions were present in the EC and amygdala, compared to vehicle-treated controls. This suggests that modulating tau pathology in the olfactory system delays its progression in interconnected CNS regions. Given that all these regions receive input from the OB, our findings reinforce the hypothesis that the olfactory pathway – particularly the OE – serves as an early site for pathology development and as a potential entry point for the spread of pathology to the PC and EC. These findings highlight the potential of intranasal drug delivery to target NDs via the olfactory pathway, offering a way to reduce the side effects associated with systemic administration [34]. Research in mouse models has demonstrated that nasal tau immunotherapy can clear intracellular tau pathology and improve cognitive function in aged tauopathy mice [25]. In humans, the intranasal route has also been explored in multiple studies as a promising therapeutic strategy for various NDs [18, 29, 40].

## Limitations of this study

In this study, the limitations of using PS19 mice, a transgenic model with random transgene insertion and artificial overexpression driven by the mouse prion protein promoter, must be taken into account when interpreting the results [53]. It is also important to emphasize that this is exclusively a tauopathy model, with no amyloid pathology present. As a result, we were unable to study the impact of co-existing amyloid pathology on the appearance and progression of tau pathology in the olfactory and central nervous systems, which would be of interest but particularly challenging to carry out. We focused on tauopathy rather than on amyloid pathology for its closer correlation to neurodegeneration. To partially address this limitation and give indication about the relevance of our results, we incorporated results obtained from human samples with a *postmortem* diagnosis of early Braak stage pathology.

## Conclusions

Our study highlights that pTau appears early in the OE of PS19 mice and subsequently spreads to the CNS following neuroanatomical pathways, like the olfactory and perforant pathways. Despite the presence of tau lesions in regions of the olfactory system, the olfactory threshold of PS19 mice was not significantly impaired. Intranasal administration of ZnSO_4_ in PS19 mice was used to mitigate the progression of tau pathology in CNS regions. The treatment significantly reduced pretangle-like tau pathology in the PC, EC, and amygdala. We additionally confirmed the appearance of pTau signal in the olfactory regions of Braak stage I/II patients. Overall, our findings support the spread of tau pathology according to anatomical neuronal connectivity, highlight the therapeutic potential of the OE region in tauopathies, and suggest its likely contribution to the progression of pathology in the CNS.

## Declarations

### Funding

NS is funded by a Chargé de Recherche postdoctoral fellowship from the F.R.S.-FNRS. This work was supported by grants from the SAO-FRA Alzheimer Research Foundation (SAO-FRA 2018/0025), UCLouvain Action de Recherche Concertée (Towards the development of new, non-invasive diagnostic tools for neurodegenerative pathologies), Fondation Louvain, Queen Elisabeth Medical Foundation (FMRE AlzHex), F.R.S.-FNRS (FNRS J.0106.22) to PKC and SAO-FRA Alzheimer Research Foundation (SAO-FRA 2020/0028) to NS. Fonds Wetenschappelijk Onderzoek Vlaanderen (FWO: G065721N, G024925N (DRT)). FWO (1225725N) (SOT).

### Disclosure of potential conflicts of interest

DRT and SOT received consultant honoraria from Muna Therapeutics. DRT collaborated with Novartis Pharma AG (Switzerland), and GE Healthcare (UK). The other authors have no competing interests to declare that are relevant to the content of this article.

### Statement on welfare

All animal procedures were conducted in accordance with institutional and European guidelines and approved by the UCLouvain Ethical Committee for Animal Welfare (2021/UCL/MD/018). Human *postmortem* brain tissue was collected in accordance with the applicable legislation in Belgium. The recruitment protocols for the collection of human brains were approved by the ethical committee (2020/02JUL/355).

### Data available statements

All datasets generated and analyzed during this study are included in this published article and its supplementary material.

### Authors contribution

Conceptualization: MD, PKC; Methodology: MD, EP, EB, CH, MB, ED; Formal analysis and investigation: MD; Writing – original draft preparation: MD; Writing – review and editing: MD, EB, NS, SOT, KL, ED, DRT, BH, PKC; Funding: AD, CH, BH, PKC; Supervision: PKC. All authors read and approved the final manuscript.

## Acknowledgements

We thank Olivier Schakman for conducting behavioral studies in mice. We thank Yasmine Salman, Benoît Lengelé and Catherine Behets for collecting and handling the human samples. We thank Marc de Bournonville and Olivier Van Kerk for their technical support. We finally thank the team of Peter Davies, who kindly provided the PHF-1 antibody.

## List of Abbreviations

ABs: Axon bundles
AD: Alzheimer’s disease
CNS: Central nervous system
DG: Dentate gyrus
EC: Entorhinal cortex
EPL: External plexiform layer
FTLD: Frontotemporal lobe degeneration
GCL: Granular cell layer
GL: Glomerular cell layer
IPL: Internal plexiform layer
MAPT: Microtubule associated protein tau
MCI: Mild cognitive impairment
MCL: Mitral cell layer
NDs: Neurodegenerative diseases
NFTs: Neurofibrillary tangles
OB: Olfactory bulb
OC: Olfactory cortex
OE: Olfactory epithelium
OI: Olfactory impairment
OMP: Olfactory mature protein
ONL: Olfactory nerve layer
OS: Olfactory system
OSNs: Olfactory sensory neurons
PC: Piriform cortex
PHF: Paired helical filaments
RE: Respiratory epithelium
WT: Wild-type

## References

1. Ahnaou, A., D. Rodriguez-Manrique, R. Biermans, S. Embrechts, N.V. Manyakov and W.H. Drinkenburg, Functional Alterations in the Olfactory Neuronal Circuit Occur before Hippocampal Plasticity Deficits in the P301S Mouse Model of Tauopathy: Implications for Early Diagnosis and Translational Research in Alzheimer’s Disease. Int J Mol Sci, 2020. 21(15). 10.3390/ijms21155431

2. Amani, M., J.C. Lauterborn, A.A. Le, B.M. Cox, W. Wang, J. Quintanilla, C.D. Cox, C.M. Gall, and G. Lynch, Rapid Aging in the Perforant Path Projections to the Rodent Dentate Gyrus. J Neurosci, 2021. 41(10): p. 2301–2312. 10.1523/jneurosci.2376-20.2021

3. Aragão Gomes, L., V. Uytterhoeven, D. Lopez-Sanmartin, S.O. Tomé, T. Tousseyn, R. Vandenberghe, M. Vandenbulcke, C.A.F. von Arnim, P. Verstreken, and D.R. Thal, Maturation of neuronal AD-tau pathology involves site-specific phosphorylation of cytoplasmic and synaptic tau preceding conformational change and fibril formation. Acta Neuropathologica, 2021. 141(2): p. 173–192. 10.1007/s00401-020-02251-6

4. Arnold, S.E., G.S. Smutzer, J.Q. Trojanowski and P.J. Moberg, Cellular and molecular neuropathology of the olfactory epithelium and central olfactory pathways in Alzheimer’s disease and schizophrenia. Ann N Y Acad Sci, 1998. 855: p. 762–75. 10.1111/j.1749-6632.1998.tb10656.x

5. Attems, J. and K.A. Jellinger, Olfactory tau pathology in Alzheimer disease and mild cognitive impairment. Clin Neuropathol, 2006. 25(6): p. 265–71.

6. Attems, J., F. Lintner and K.A. Jellinger, Olfactory involvement in aging and Alzheimer’s disease: an autopsy study. J Alzheimers Dis, 2005. 7(2): p. 149–57; discussion 173-80. 10.3233/jad-2005-7208

7. Attems Johannes, W.L., and Jellinger Kurt A., Olfactory bulb involvement in neurodegenerative diseases. Acta Neuropathologica, 2014. 127: p. 459–475.

8. Barrios, A.W., G. Núñez, P. Sánchez Quinteiro and I. Salazar, Anatomy, histochemistry, and immunohistochemistry of the olfactory subsystems in mice. Frontiers in Neuroanatomy, 2014. 8. 10.3389/fnana.2014.00063

9. Bathini, P., E. Brai and L.A. Auber, Olfactory dysfunction in the pathophysiological continuum of dementia. Ageing Res Rev, 2019. 55: p. 100956. 10.1016/j.arr.2019.100956

10. Braak, H., I. Alafuzoff, T. Arzberger, H. Kretzschmar and K. Del Tredici, Staging of Alzheimer disease-associated neurofibrillary pathology using paraffin sections and immunocytochemistry. Acta Neuropathol, 2006. 112(4): p. 389–404. 10.1007/s00401-006-0127-z

11. Braak, H. and E. Braak, Neuropathological stageing of Alzheimer-related changes. Acta Neuropathol. (Berl), 1991. 82(4): p. 239–259.

12. Chen, C.R., C. Kachramanoglou, D. Li, P. Andrews and D. Choi, Anatomy and cellular constituents of the human olfactory mucosa: a review. J Neurol Surg B Skull Base, 2014. 75(5): p. 293–300. 10.1055/s-0033-1361837

13. Clavaguera, F., T. Bolmont, R.A. Crowther, D. Abramowski, S. Frank, A. Probst, G. Fraser, A.K. Stalder, M. Beibel, M. Staufenbiel, M. Jucker, M. Goedert, and M. Tolnay, Transmission and spreading of tauopathy in transgenic mouse brain. Nat Cell Biol, 2009. 11(7): p. 909–13. 10.1038/ncb1901

14. Dan, X., N. Wechter, S. Gray, J.G. Mohanty, D.L. Croteau and V.A. Bohr, Olfactory dysfunction in aging and neurodegenerative diseases. Ageing Res Rev, 2021. 70: p. 101416. 10.1016/j.arr.2021.101416

15. DeTure, M.A. and D.W. Dickson, The neuropathological diagnosis of Alzheimer’s disease. Molecular Neurodegeneration, 2019. 14(1): p. 32. 10.1186/s13024-019-0333-5

16. Devi, G., Handbook of Clinical Neurology, Chapter 12 – The tauopathies Vol. 196. 2023.

17. Diez, I., L. Ortiz-Terán, T.S.C. Ng, M.W. Albers, G. Marshall, W. Orwig, C.-m. Kim, E. Bueichekú, V. Montal, J. Olofsson, P. Vannini, G. El Fahkri, R. Sperling, K. Johnson, H.I.L. Jacobs, and J. Sepulcre, Tau propagation in the brain olfactory circuits is associated with smell perception changes in aging. Nature Communications, 2024. 15(1): p. 4809. 10.1038/s41467-024-48462-3

18. Djupesland, P.G., J.C. Messina and R.A. Mahmoud, The Nasal Approach to Delivering Treatment for Brain Diseases: An Anatomic, Physiologic, and Delivery Technology Overview. Therapeutic Delivery, 2014. 5(6): p. 709–733. 10.4155/tde.14.41

19. Dorman, D.C., 13.15 – Olfactory System, in Comprehensive Toxicology (Second Edition), C.A. McQueen, Editor. 2010, Elsevier: Oxford. p. 263–276.

20. Doty, R.L., Olfactory dysfunction in neurodegenerative diseases: is there a common pathological substrate? Lancet Neurology, 2017. 16(6): p. 478–488.

21. Dujardin, S. and B.T. Hyman, Tau Prion-Like Propagation: State of the Art and Current Challenges. Adv Exp Med Biol, 2019. 1184: p. 305–325. 10.1007/978-981-32-9358-8_23

22. Fatuzzo, I., G.F. Niccolini, F. Zoccali, L. Cavalcanti, M.G. Bellizzi, G. Riccardi, M. de Vincentiis, M. Fiore, C. Petrella, A. Minni, and C. Barbato, *Neurons, Nose,* and Neurodegenerative Diseases: Olfactory Function and Cognitive Impairment. Int J Mol Sci, 2023. 24(3). 10.3390/ijms24032117

23. Franco, R., C. Garrigós and J. Lillo, The Olfactory Trail of Neurodegenerative Diseases. Cells, 2024. 13(7). 10.3390/cells13070615

24. Fujishiro, H., Y. Tsuboi, W.L. Lin, H. Uchikado and D.W. Dickson, Co-localization of tau and alpha-synuclein in the olfactory bulb in Alzheimer’s disease with amygdala Lewy bodies. Acta Neuropathol, 2008. 116(1): p. 17–24. 10.1007/s00401-008-0383-1

25. Gaikwad, S., N. Puangmalai, M. Sonawane, M. Montalbano, R. Price, M.S. Iyer, A. Ray, S. Moreno, and R. Kayed, Nasal tau immunotherapy clears intracellular tau pathology and improves cognitive functions in aged tauopathy mice. Sci Transl Med, 2024. 16(754): p. eadj5958. 10.1126/scitranslmed.adj5958

26. Gallyas, F., Silver staining of Alzheimer’s neurofibrillary changes by means of physical development. Acta Morphol Acad Sci Hung, 1971. 19(1): p. 1–8.

27. Gao, Y.L., N. Wang, F.R. Sun, X.P. Cao, W. Zhang and J.T. Yu, Tau in neurodegenerative disease. Ann Transl Med, 2018. 6(10): p. 175. 10.21037/atm.2018.04.23

28. Greenberg, S.G., P. Davies, J.D. Schein and L.I. Binder, Hydrofluoric acid-treated tau PHF proteins display the same biochemical properties as normal tau. J Biol Chem, 1992. 267(1): p. 564–9.

29. Guo, H., G. Wang, Z. Zhai, J. Huang, Z. Huang, Y. Zhou, X. Xia, Z. Yao, Y. Huang, Z. Zhao, C. Wu, and X. Zhang, Rivastigmine nasal spray for the treatment of Alzheimer’s Disease: Olfactory deposition and brain delivery. International Journal of Pharmaceutics, 2024. 652: p. 123809. 10.1016/j.ijpharm.2024.123809

30. Hannsjörg Schröder, N.M., Stefan Huggenberger, Neuroanatomy of the Mouse. 2020: Springer Cham. 350.

31. Harding, J.W., T.V. Getchell and F.L. Margolis, Denervation of the primary olfactory pathway in mice. V. Long-term effect of intranasal ZnSO4 irrigation on behavior, biochemistry and morphology. Brain Res, 1978. 140(2): p. 271–85. 10.1016/0006-8993(78)90460-2

32. Keller, M., Q. Douhard, M.J. Baum and J. Bakker, Destruction of the main olfactory epithelium reduces female sexual behavior and olfactory investigation in female mice. Chem Senses, 2006. 31(4): p. 315–23. 10.1093/chemse/bjj035

33. Kondo, K., S. Kikuta, R. Ueha, K. Suzukawa and T. Yamasoba, Age-Related Olfactory Dysfunction: Epidemiology, Pathophysiology, and Clinical Management. Front Aging Neurosci, 2020. 12: p. 208. 10.3389/fnagi.2020.00208

34. Kovács, T., The olfactory system in Alzheimer’s disease: Pathology, pathophysiology and pathway for therapy. Translational Neuroscience, 2013. 4(1): p. 34–45. doi: 10.2478/s13380-013-0108-3

35. Kovács, T., N.J. Cairns and P.L. Lantos, Olfactory centres in Alzheimer’s disease: olfactory bulb is involved in early Braak’s stages. NeuroReport, 2001. 12(2): p. 285–288.

36. Lad, M., W. Sedley and T.D. Griffiths, Sensory Loss and Risk of Dementia. Neuroscientist, 2024. 30(2): p. 247–259. 10.1177/10738584221126090

37. LaFever, B.J. and F. Imamura, Effects of nasal inflammation on the olfactory bulb. J Neuroinflammation, 2022. 19(1): p. 294. 10.1186/s12974-022-02657-x

38. Laska, M., Human and Animal Olfactory Capabilities Compared. 2017. p. 675–689.

39. Li, S., W. Li, X. Wu, J. Li, J. Yang, C. Tu, X. Ye, and S. Ling, Olfactory deficit is associated with mitral cell dysfunction in the olfactory bulb of P301S tau transgenic mice. Brain Res Bull, 2019. 148: p. 34–45. 10.1016/j.brainresbull.2019.03.006

40. Lofts, A., F. Abu-Hijleh, N. Rigg, R.K. Mishra and T. Hoare, Using the Intranasal Route to Administer Drugs to Treat Neurological and Psychiatric Illnesses: Rationale, Successes, and Future Needs. CNS Drugs, 2022. 36(7): p. 739–770. 10.1007/s40263-022-00930-4

41. Lundström, J.N., S. Boesveldt and J. Albrecht, Central Processing of the Chemical Senses: an Overview. ACS Chem Neurosci, 2011. 2(1): p. 5–16. 10.1021/cn1000843

42. Martel, G., A. Simon, S. Nocera, S. Kalainathan, L. Pidoux, D. Blum, S. Leclère-Turbant, J. Diaz, D. Geny, E. Moyse, C. Videau, L. Buée, J. Epelbaum, and C. Viollet, Aging, but not tau pathology, impacts olfactory performances and somatostatin systems in THY-Tau22 mice. Neurobiol Aging, 2015. 36(2): p. 1013–28. 10.1016/j.neurobiolaging.2014.10.033

43. Matulionis, D.H., Ultrastructural study of mouse olfactory epithelium following destruction by ZnSO4 and its subsequent regeneration. Am J Anat, 1975. 142(1): p. 67–89. 10.1002/aja.1001420106

44. McBride, K., B. Slotnick and F.L. Margolis, Does intranasal application of zinc sulfate produce anosmia in the mouse? An olfactometric and anatomical study. Chem Senses, 2003. 28(8): p. 659–70. 10.1093/chemse/bjg053

45. Noto, T., G. Zhou, Q. Yang, G. Lane and C. Zelano, Human Primary Olfactory Amygdala Subregions Form Distinct Functional Networks, Suggesting Distinct Olfactory Functions. Frontiers in Systems Neuroscience, 2021. 15. 10.3389/fnsys.2021.752320

46. Ohm, T.G. and H. Braak, Olfactory bulb changes in Alzheimer’s disease. Acta Neuropathologica, 1987. 73(4): p. 365–369. 10.1007/BF00688261

47. Orr, M.E., A.C. Sullivan and B. Frost, A Brief Overview of Tauopathy: Causes, Consequences, and Therapeutic Strategies. Trends Pharmacol Sci, 2017. 38(7): p. 637–648. 10.1016/j.tips.2017.03.011

48. Otvos, L., Jr., L. Feiner, E. Lang, G.I. Szendrei, M. Goedert and V.M. Lee, Monoclonal antibody PHF-1 recognizes tau protein phosphorylated at serine residues 396 and 404. J Neurosci Res, 1994. 39(6): p. 669–73. 10.1002/jnr.490390607

49. Parvand, M. and C.H. Rankin, Is There a Shared Etiology of Olfactory Impairments in Normal Aging and Neurodegenerative Disease? J Alzheimers Dis, 2020. 73(1): p. 1–21. 10.3233/jad-190636

50. Poulin, S.P., R. Dautoff, J.C. Morris, L.F. Barrett and B.C. Dickerson, Amygdala atrophy is prominent in early Alzheimer’s disease and relates to symptom severity. Psychiatry Res, 2011. 194(1): p. 7–13. 10.1016/j.pscychresns.2011.06.014

51. Ramirez, D.M.O., J.D. Whitesell, N. Bhagwat, T.L. Thomas, A.D. Ajay, A. Nawaby, B. Delatour, S. Bay, P. LaFaye, J.E. Knox, J.A. Harris, J.P. Meeks, and M.I. Diamond, Endogenous pathology in tauopathy mice progresses via brain networks. bioRxiv, 2023. 10.1101/2023.05.23.541792

52. Rey, N.L., D.W. Wesson and P. Brundin, The olfactory bulb as the entry site for prion-like propagation in neurodegenerative diseases. Neurobiology of Disease, 2018. 109: p. 226–248. 10.1016/j.nbd.2016.12.013

53. Sahara, N. and R. Yanai, Limitations of human tau-expressing mouse models and novel approaches of mouse modeling for tauopathy. Frontiers in Neuroscience, 2023. 17. 10.3389/fnins.2023.1149761

54. Samudra, N., C. Lane-Donovan, L. VandeVrede and A.L. Boxer, Tau pathology in neurodegenerative disease: disease mechanisms and therapeutic avenues. J Clin Invest, 2023. 133(12). 10.1172/jci168553

55. Shimizu, S., I. Tojima, K. Nakamura, H. Kouzaki, T. Kanesaka, N. Ogawa, Y. Hashizume, H. Akatsu, A. Hori, I. Tooyama, and T. Shimizu, A Histochemical Analysis of Neurofibrillary Tangles in Olfactory Epithelium, a Study Based on an Autopsy Case of Juvenile Alzheimer’s Disease. Acta Histochem Cytochem, 2022. 55(3): p. 93–98. 10.1267/ahc.22-00048

56. Smith, T.D. and K.P. Bhatnagar, Anatomy of the olfactory system. Handb Clin Neurol, 2019. 164: p. 17–28. 10.1016/b978-0-444-63855-7.00002-2

57. Son, G., A. Jahanshahi, S.J. Yoo, J.T. Boonstra, D.A. Hopkins, H.W.M. Steinbusch, and C. Moon, Olfactory neuropathology in Alzheimer’s disease: a sign of ongoing neurodegeneration. BMB Rep, 2021. 54(6): p. 295–304. 10.5483/BMBRep.2021.54.6.055

58. Sun, Y., Y. Guo, X. Feng, M. Jia, N. Ai, Y. Dong, Y. Zheng, L. Fu, B. Yu, H. Zhang, J. Wu, X. Yu, H. Wu, and W. Kong, The behavioural and neuropathologic sexual dimorphism and absence of MIP-3α in tau P301S mouse model of Alzheimer’s disease. Journal of Neuroinflammation, 2020. 17(1): p. 72. 10.1186/s12974-020-01749-w

59. Takehara-Nishiuchi, K., Entorhinal cortex and consolidated memory. Neuroscience Research, 2014. 84: p. 27–33. 10.1016/j.neures.2014.02.012

60. Talamo, B.R., W.H. Feng, M. Perez-Cruet, L. Adelman, K. Kosik, M.Y. Lee, L.C. Cork, and J.S. Kauer, Pathologic changes in olfactory neurons in Alzheimer’s disease. Ann N Y Acad Sci, 1991. 640: p. 1–7. 10.1111/j.1749-6632.1991.tb00182.x

61. Tzeng, W.-Y., K. Figarella and O. Garaschuk, Olfactory impairment in men and mice related to aging and amyloid-induced pathology. Pflügers Archiv – European Journal of Physiology, 2021. 473(5): p. 805–821. 10.1007/s00424-021-02527-0

62. Ubeda-Bañon, I., D. Saiz-Sanchez, A. Flores-Cuadrado, E. Rioja-Corroto, M. Gonzalez-Rodriguez, S. Villar-Conde, V. Astillero-Lopez, J.P. Cabello-de la Rosa, M.J. Gallardo-Alcañiz, J. Vaamonde-Gamo, F. Relea-Calatayud, L. Gonzalez-Lopez, A. Mohedano-Moriano, A. Rabano, and A. Martinez-Marcos, The human olfactory system in two proteinopathies: Alzheimer’s and Parkinson’s diseases. Transl Neurodegener, 2020. 9(1): p. 22. 10.1186/s40035-020-00200-7

63. Walkiewicz, G., A. Ronisz, S. Ospitalieri, G. Tsaka, S.O. Tomé, R. Vandenberghe, C.A.F. von Arnim, F. Rousseau, J. Schymkowitz, L. De Groef, and D.R. Thal, pTau pathology in the retina of TAU58 mice: association with ganglion cell degeneration and implications on seeding and propagation of pTau from human brain lysates. Acta Neuropathologica Communications, 2024. 12(1): p. 194. 10.1186/s40478-024-01907-8

64. Witter, M.P., P.A. Naber, T. van Haeften, W.C.M. Machielsen, S.A.R.B. Rombouts, F. Barkhof, P. Scheltens, and F.H. Lopes da Silva, Cortico-hippocampal communication by way of parallel parahippocampal-subicular pathways. Hippocampus, 2000. 10(4): p. 398–410. 10.1002/1098-1063(2000)10:4<398::AID-HIPO6>3.0.CO;2-K

65. Yan, Y., A. Aierken, C. Wang, D. Song, J. Ni, Z. Wang, Z. Quan, and H. Qing, A potential biomarker of preclinical Alzheimer’s disease: The olfactory dysfunction and its pathogenesis-based neural circuitry impairments. Neurosci Biobehav Rev, 2022. 132: p. 857–869. 10.1016/j.neubiorev.2021.11.009

66. Yang, S., W.L. Kuan and M.G. Spillantini, Progressive tauopathy in P301S tau transgenic mice is associated with a functional deficit of the olfactory system. Eur J Neurosci, 2016. 44(6): p. 2396–403. 10.1111/ejn.13333

67. Yasumasa Yoshiyama, M.H., Bin Zhang, Shu-Ming Huang, Nobuhisa Iwata, Takaomi C. Saido, Jun Maeda, Tetsuya Suhara, John Q. Trojanowski, and Virginia M-Y Lee, Synapse loss and microglial activation precede tangles in a P301S tauopathy mouse model. Neuron, 2007. 53(3): p. 337–351. 10.1016/j.neuron.2007.01.010

68. Yu, H., F. Wang, D. Jia, S. Bi, J. Gong, J.J. Wu, Y. Mao, J. Chen, and G.S. Chai, Pathological features and molecular signatures of early olfactory dysfunction in 3xTg-AD model mice. CNS Neurosci Ther, 2024. 30(2): p. e14632. 10.1111/cns.14632

69. Zhang, J., Z. Zhao, S. Sun, J. Li, Y. Wang, J. Dong, S. Yang, Y. Lou, J. Yang, W. Li, and S. Li, Olfactory Evaluation in Alzheimer’s Disease Model Mice. Brain Sci, 2022. 12(5). 10.3390/brainsci12050607

70. Zhou, X., P. Kumar, D.J. Bhuyan, S.O. Jensen, T.L. Roberts and G.W. Münch, Neuroinflammation in Alzheimer’s Disease: A Potential Role of Nose-Picking in Pathogen Entry via the Olfactory System? Biomolecules, 2023. 13(11). 10.3390/biom13111568

