## Supplementary information for "The Olfactory Epithelium: A Critical Gateway for Pathological Tau Propagation and a Target for Mitigating Tauopathy in the Central Nervous System"

**Supplementary methods**

**Protein Extraction**

Proteins were isolated from snap-frozen samples of OE, OB, and hippocampus. Lysis buffer (LB) containing 50 mM Tris base (pH 7.5), 150 mM NaCl, 2 mM EDTA, 1% NP-40, % glycerol was prepared with an adjusted pH of 8.0. Before each experiment phosphatase inhibitor (Merck) and protease inhibitor cocktail (Roche) were added in the lysis buffer. A specific amount of ice-cold LB was added to each sample: 50 μL for the pooled right and left OE, 150 µL for a single OB and 200 µL for a single hippocampus.

Tissues were homogenized by sonication for 15 seconds, and then centrifuged at 16.100 g for 15 minutes at 4°C. The supernatants containing protein extracts were collected and stored at -80°C. Protein concentration was quantified using Pierce™ BCA Protein assay kit (Invitrogen, Thermo Fisher Scientific). Each sample was diluted 20 times for dosage.

**Western Blot**

A total of 20 μg of protein extracts were mixed in a final volume of 20 μL with NuPAGE™ LDS Sample Buffer (Invitrogen, Thermo Fisher Scientific) and 50 mM dithiothreitol (DTT). The samples were vortexed, spun down and heated at 70°C for 10 minutes. Samples were loaded into polyacrylamide gel (NuPAGETM 4-12% Bis Tris Gel, InvitrogenTM #NP0336BOX) using SeeBlue™ Plus2 pre-stained (Thermo Fisher Scientific) as standard. Electrophoresis was run for 3 hours at 110 volts. This step was followed by protein transfer onto a nitrocellulose membrane (Thermo Fisher Scientific) for 2 hours at 30 volts. Membranes were blocked with 5% non-fat powdered milk in PBS-Tween 0,1% (PBS-T) for one hour with agitation, then rinsed and incubated overnight at 4°C with the primary antibodies (supplementary table S1) of interest prepared in PBS-T.

The next day, membranes were rinsed twice in PBS-T for 10 minutes and incubated with horseradish peroxidase (HRP)-conjugated species-specific secondary antibodies (Sigma-Aldrich) for one hour at room temperature under agitation. After a final rinse, the signal was detected by incubating with chemiluminescence ECL substrate (1:1 reagent A and B) for 2 minutes. Membranes were then exposed to chemiluminescence film and developed. ImageJ software was used to quantify protein band intensity, normalized to actin intensity.

**Masson’s trichrome stain**

Paraffin sections (10μm) were processed through a series of solutions for deparaffinization and staining. Sections were immersed sequentially: three times for 5 minutes in toluene, followed by 3x5 minutes in absolute isopropanol, 95% isopropanol, and 70% isopropanol. After a 10-minute rinse in running water and a 5-minute rinse in distilled water, the sections were incubated in Mayer’s hemalum solution for 10 minutes. This was followed by identical rinsing steps. The sections were stained in ponceau fuchsine solution for 3 minutes, rinsed in distilled water for 1 minute, and incubated in 1% phosphomolybdic acid for 5 minutes before a quick rinse in distilled water. A final staining step with 1% aniline blue for 3 minutes was performed, followed by two distilled water rinses.

The sections were dehydrated through a graded series of isopropanol (70%, absolute x2), followed by 2x5 minutes in toluene and 5 minutes in xylene. The sections were mounted using DPX mounting medium (Sigma-Aldrich, #1.00579.0500) and left to dry overnight at room temperature. Images were captured using a Zeiss Axioskop 40 microscope.

**Supplementary Figures and Legends**

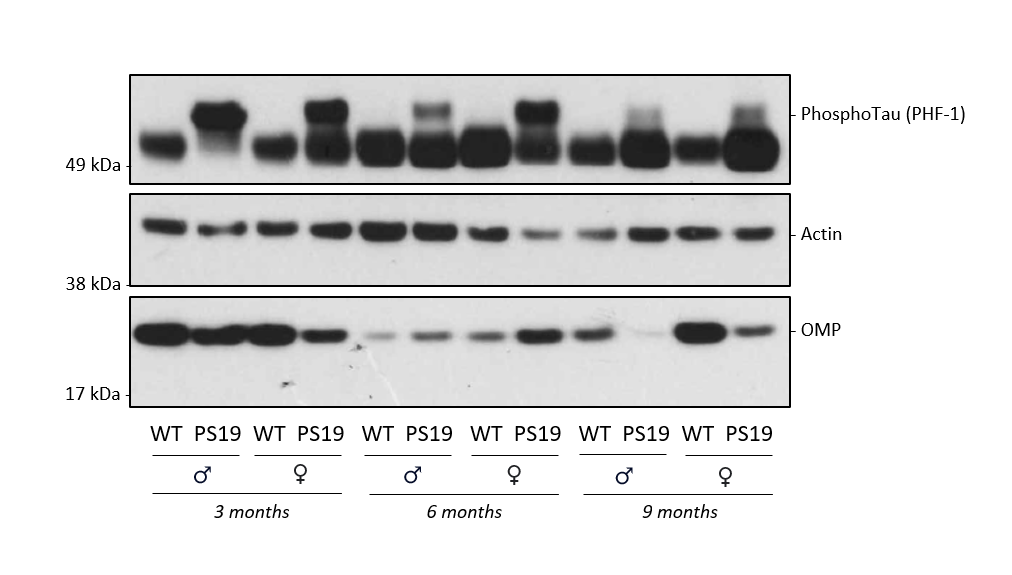

**Fig. S1 pTau expression in the OE of PS19 mice.** pTau (**PHF-1** (Ser396, Ser404)) is detected in the OE of PS19 or WT males and females at 3, 6 and 9 months. pTau detected at around 60kDa using this antibody is present only in PS19 mice, as soon as 3 months of age, both in males and females. No signal around 60kDa is observed in the WT mice regardless of ageing stage. The **OMP** antibody was used to recognize the olfactory mature protein expressed by mature OSNs (19kDa).

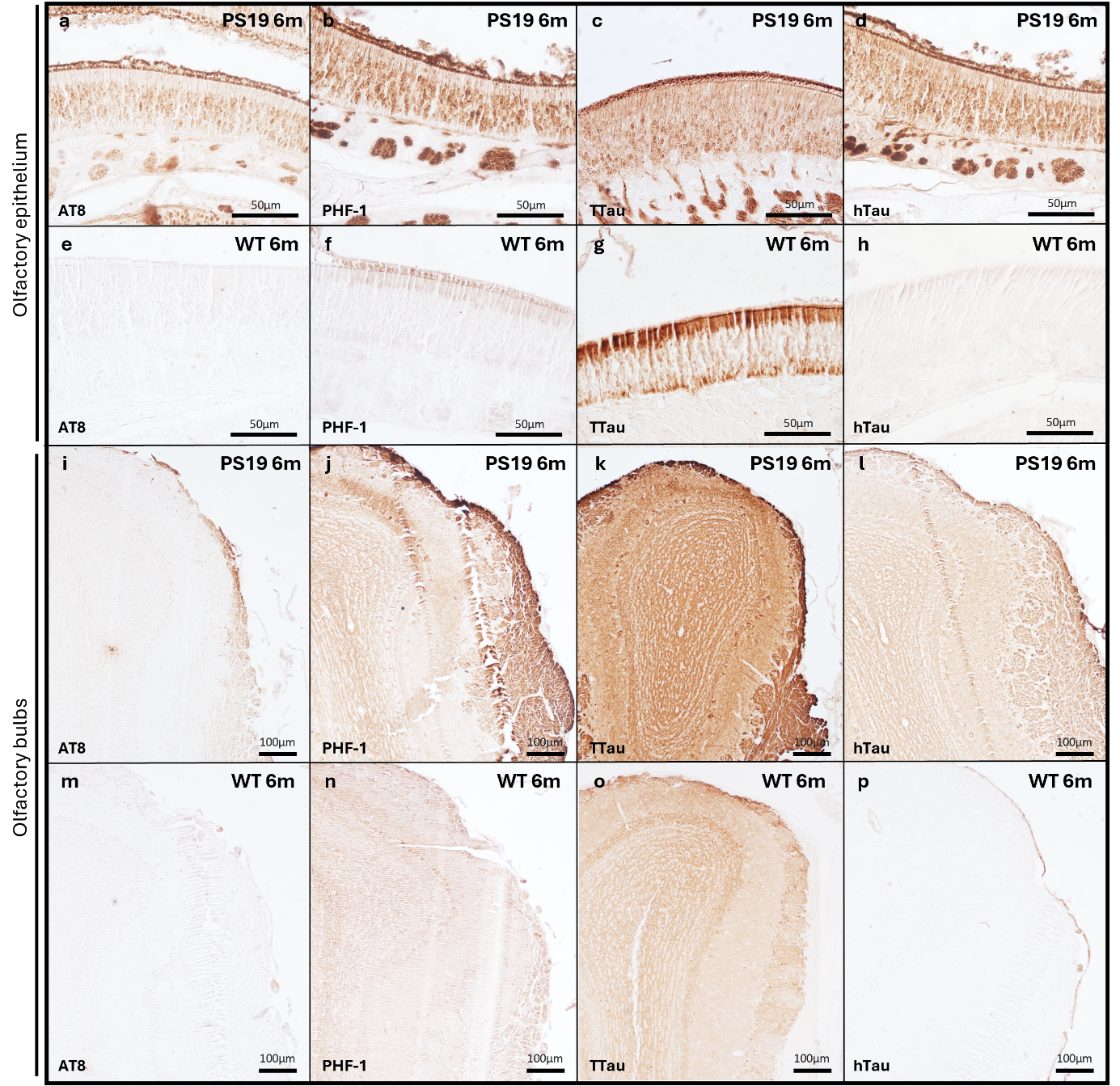

**Fig. S2 Tau protein expression in the OE and OB of PS19 and WT mice.** The OE and OB from 6-month-old PS19 and WT mice were immunohistochemically stained with **AT8** (a,e,i,m), **PHF-1** (b,f,j,n), **TTau** (c,g,k,o) and **hTau** (d,h,l,p). pTau (**AT8**) is observed in the middle stratum and ABs of the OE of PS19 mice (a), while no **AT8** signal is detected in the OE of WT mice (e). In the OB, **AT8** staining is present in the ONL and MCL of PS19 mice (i), whereas no AT8 signal is observed in WT mice (m). **PHF-1** staining reveals pTau in the middle stratum and ABs of the OE of PS19 mice (b), with some signal also detected in the apical region of the OE of WT mice (f). In the OB, **PHF-1** staining is present in the ONL and MCL of both PS19 (j) and WT (n) mice. Total tau (**TTau**) is detected throughout all strata of the OE in PS19 mice (c), whereas in WT mice, **TTau** staining is restricted to the basal and apical regions of the OE (g). In the OB, **TTau** staining is observed in the ONL and MCL of PS19 mice (k) but is limited to the MCL in WT mice (o). Like the **AT8** signal, **hTau** immunoreactivity is observed in the middle stratum and ABs of the OE of PS19 mice (d), while no **hTau** signal is detected in the OE of WT mice (h). In the OB, **hTau** staining is present in the ONL and MCL of PS19 mice (l) but is absent in WT mice (p). All mice used for immunostaining were **males**.

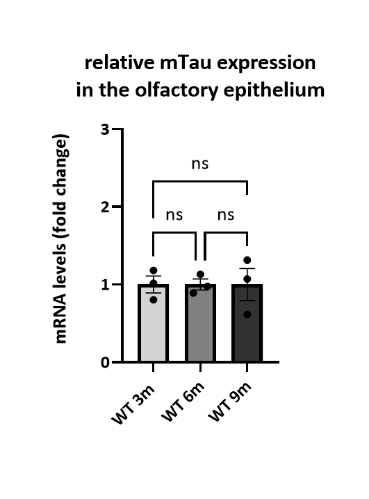

**Fig. S3 Endogenous Tau expression in the OE of WT mice.** mRNA levels of endogenous murine Tau (mTau) were measured by RT-qPCR in OE extracts from WT and PS19 mice at 3, 6 and 9 months (One-way ANOVA, n=3). Endogenous Tau protein expression in the OE is confirmed and is stable over time in WT mice.

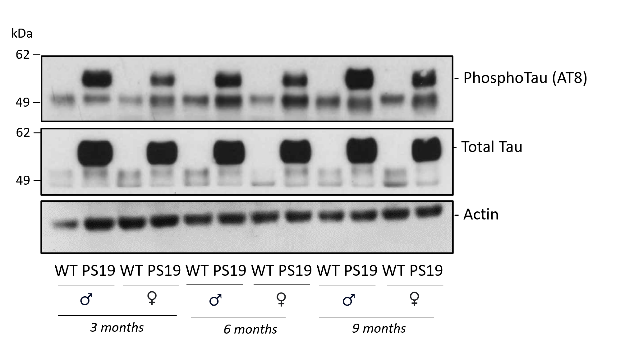

**Fig. S4 pTau expression in the OB of PS19 mice.** pTau (**AT8**) and **Total Tau** are detected around 60kDa in the OB of PS19 males and females at 3, 6 and 9 months. This upper band is not present in the WT mice regardless of ageing stage. Actin was used as loading control.

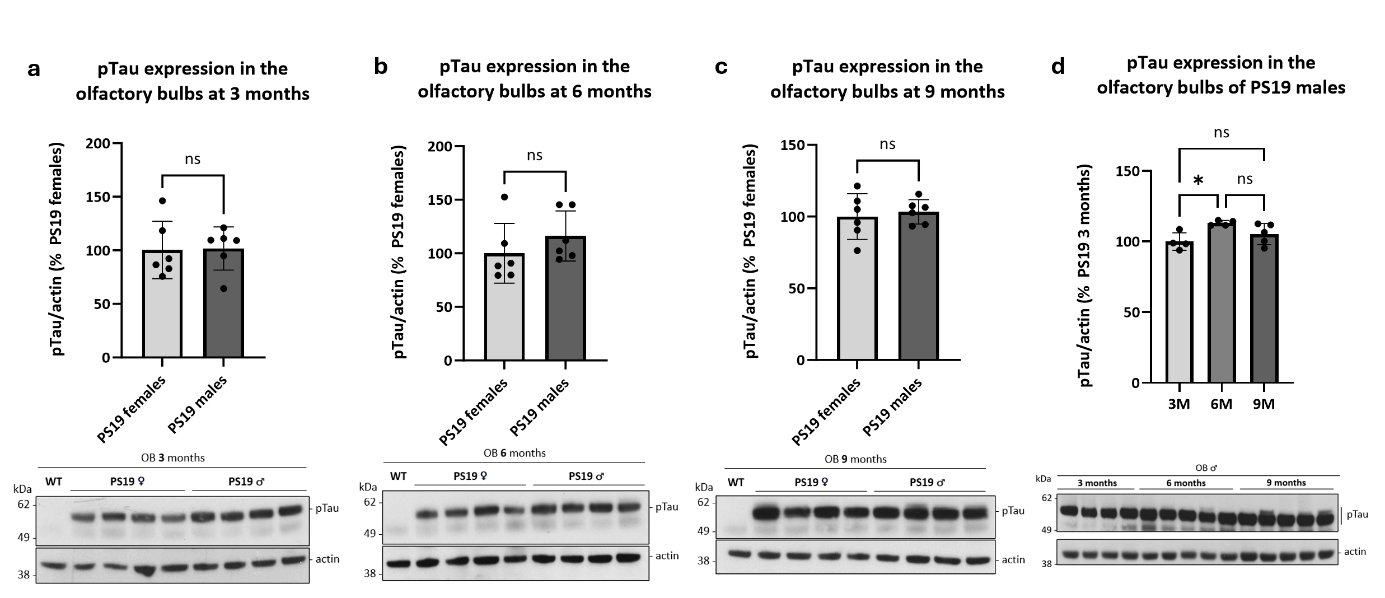

**Fig. S5 pTau expression in the OB of PS19 mice.** Western blot analyses showing pTau (**PHF-1**) relative protein levels in the OB of PS19 mice at 3, 6 and 9 months (a-d). Actin was used as loading control, and the levels in the females’ group (a-c) or at 3 months (d) were set as 100%. It shows no significant difference in pTau expression between males and females (a-c) (One-way ANOVA with Tukey’s post-hoc analysis, n = 6 mice/group). pTau expression is shown as stable between 3 and 9 months in the OB (d). *P < 0.05 (One-way ANOVA with Tukey’s post-hoc analysis, n = 4-5 mice/group).

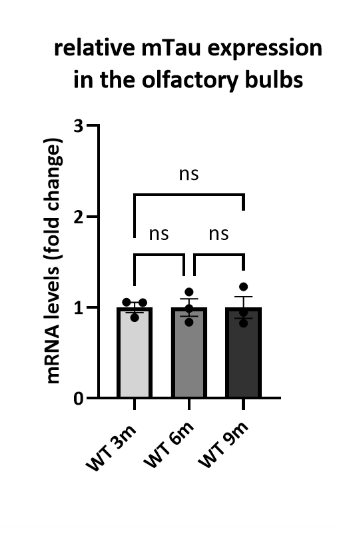

**Fig. S6 Endogenous Tau expression in the OB of WT mice.** mRNA levels of endogenous mTau expression were measured by RT-qPCR in OB extracts from WT mice at 3, 6 and 9 months (One-way ANOVA, n=3). Endogenous Tau protein expression in the OB is confirmed and is stable over time in WT mice.

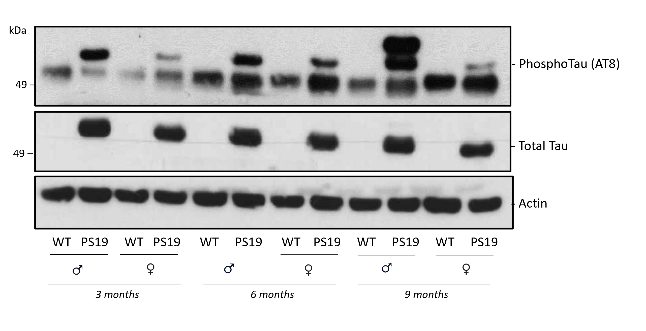

**Figure S7 pTau expression in the hippocampus of PS19 mice.** pTau (**AT8**) and **Total Tau** are detected around 60kDa in the OB of PS19 males and females at 3, 6 and 9 months. This upper band is not present in the WT mice regardless of ageing stage. Actin was used as loading control.

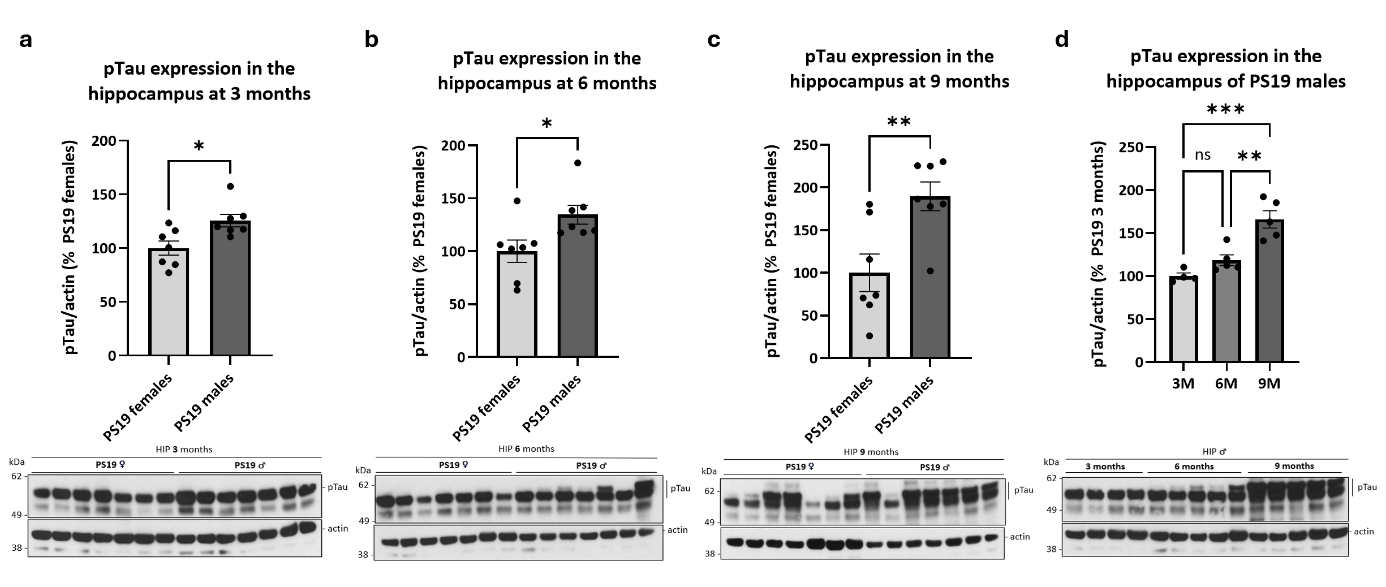

**Fig. S8 pTau expression in the hippocampus of PS19 mice.** Western blot analyses showing pTau (**PHF-1**) relative protein levels in the hippocampus of PS19 mice at 3, 6 and 9 months (a-d). Actin was used as loading control, and the levels in the females’ group (a-c) or at 3 months (d) were set as 100%. It shows a significant increase in pTau expression in males compare to females at 3, 6 and 9 months (a-c). *P < 0.05, **P < 0.01 (One-way ANOVA with Tukey’s post-hoc analysis, n = 7 mice/group). pTau expression increases significantly with age in PS19 males (d). **P < 0.01, ***P < 0.001 (One-way ANOVA with Tukey’s post-hoc analysis, n = 4-5 mice/group).

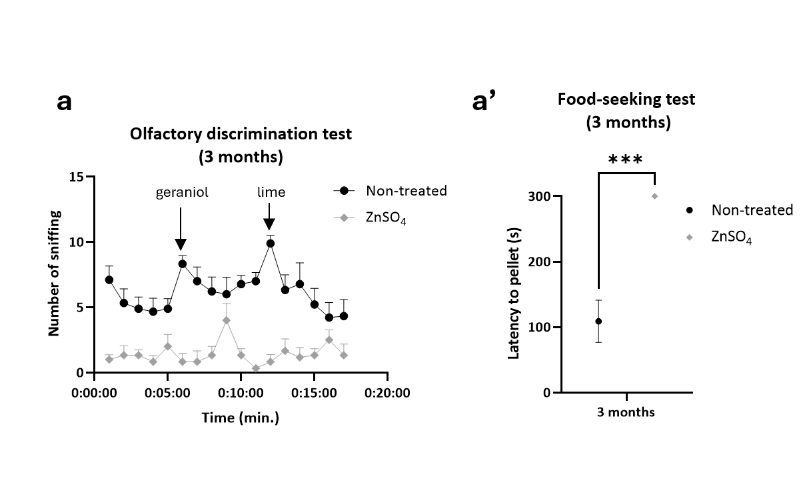

**Fig. S9 Olfactory function assessment in ZnSO_4_-treated mice.** The olfactory discrimination (a) and food-seeking tests (a’) show a significant effect (***p<0,001) of ZnSO_4_ treatment on olfactory function. For ZnSO_4_ mice; n=6 (3 months) or non-treated controls; n=9 (3 months). Data were analyzed using two-way ANOVA with Šídák’s multiple comparisons test (olfactory discrimination test) or using independent t-test (food-seeking test).

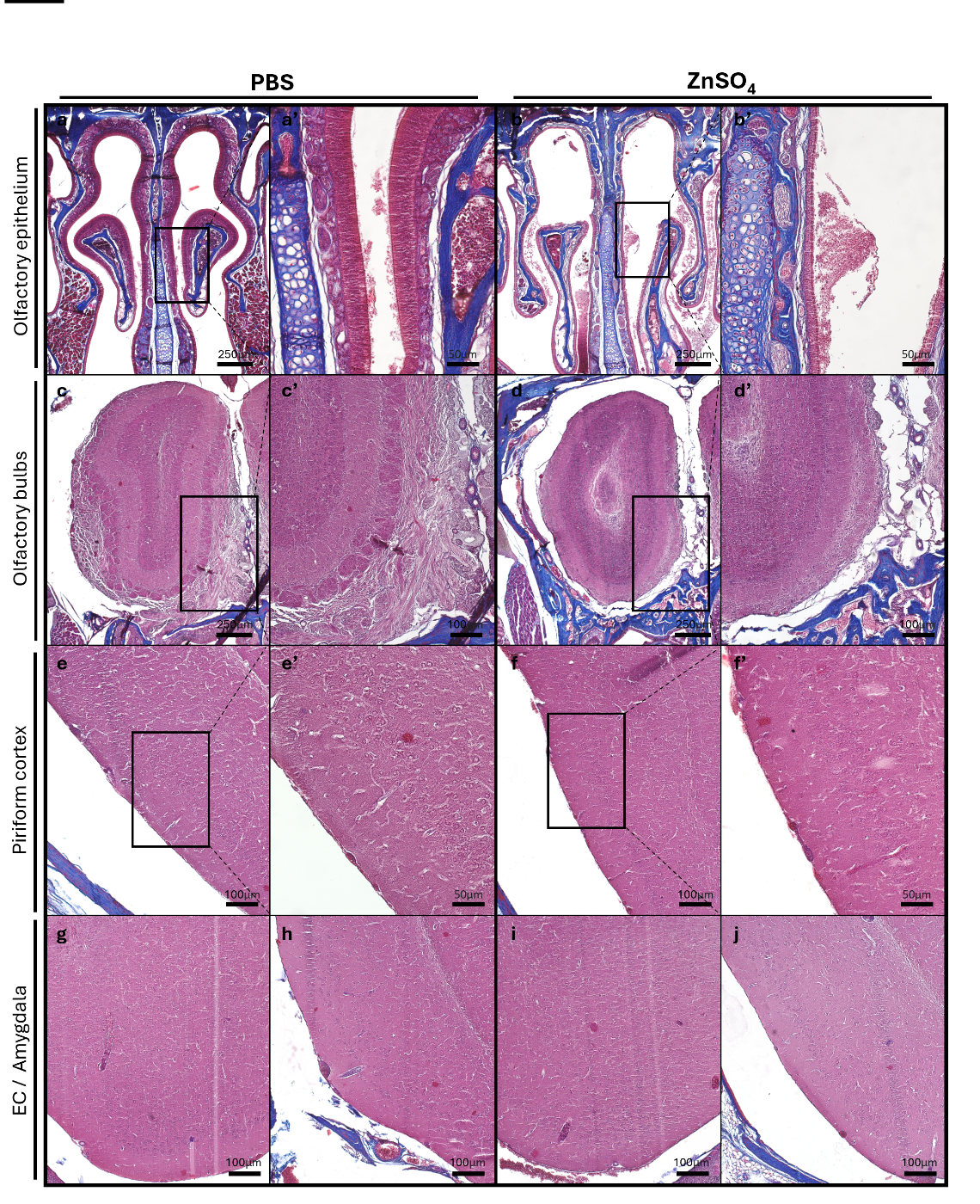

**Fig. S10 The OE, OB, PC amygdala, and EC stained with Masson’s trichrome blue following PBS or ZnSO_4_ nasal irrigation.** The staining shows that ZnSO_4_ nasal irrigation induces the stripping of the OE (b,b’) and of the ONL in the OB (d,d’) but no morphology alterations are observed in the PC (f,f’), amygdala (i) and EC (j) of 6-month-old PS19 males.

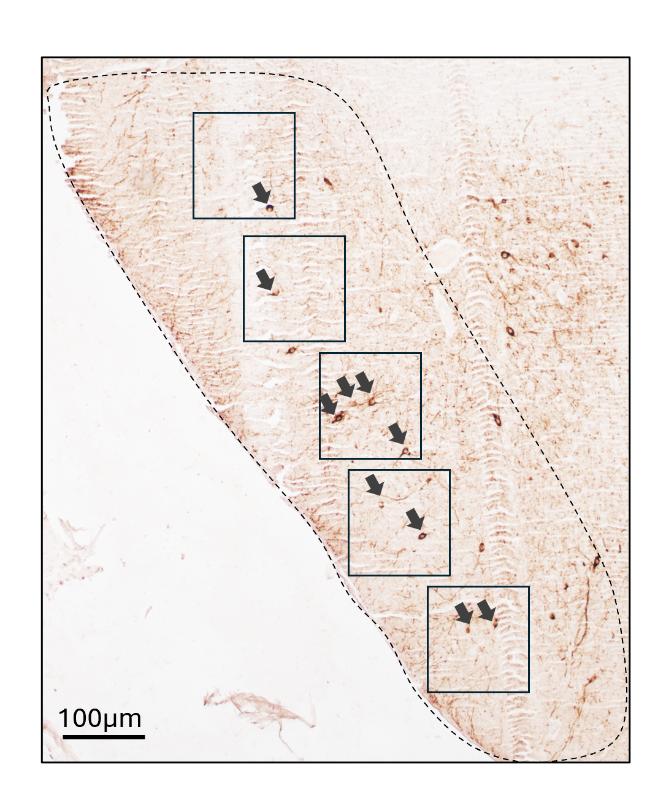

**Fig. S11 DAB quantification of the NFT-like tau accumulations in the PC.** For each region of interest, the numbers of NFT-like tau accumulations (indicated by black arrows) were counted in an equivalent area corresponding to the sum of 5 squares placed in that region. The number of NFT-like tau accumulations per mm^2^ in the vehicle and ZnSO_4_-treated conditions was compared and analyzed for each region using a Mann-Whitney test.

**Supplementary Tables**

| **Antibodies table** | | | |
| --- | --- | --- | --- |
| *Primary antibodies* | | | |
| **Antibody** | **Species** | **Dilution** | **Company (Catalog#)** |
| AT8  (Phospho-Tau (Ser202, Thr205)) | Monoclonal IgG1 Mouse | 1/250 | ThermoFisher (#MN1020) |
| Tau Total (D1M9X) | Monoclonal Rabbit | 1/1000 | Cell Signaling Technology (#46687) |
| PHF-1  *conditioned medium*  (Phospho-Tau (Ser396, Ser404)) | Monoclonal IgG1 Mouse | 1/100-200 | Kindly provided by Peter Davies’ Team |
| Tau-13 | Monoclonal IgG1 Mouse | 1/1000 | Biolegend (#835201) |
| OMP | Monoclonal Mouse | 1/50 | Santa cruz biotechnology (#sc-365818) |
| OMP | Polyclonal Goat | 1/50 | Novus biotechnology (#NB110-74751) |
| *Secondary antibodies* | | | |
| Anti-mouse biotin-labeled | IgG Goat | 1/200 | PerkinElmer, (#NEF823001) |
| Anti-Rabbit biotin-labeled | IgG Goat | 1/100 | Thermofisher (#65-6140) |
| Goat anti-Mouse IgG1 Cross Adsorbed Alexa Fluor^TM^ 647 | Goat | 1/500 | InVitrogen (#A21240) |
| Goat anti-Mouse IgG2a Cross Adsorbed Alexa Fluor™ 488 | Goat | 1/500 | InVitrogen (#A21131) |
| Goat anti-Rabbit IgG (H+L) Highly Cross-Adsorbed Alexa Fluor^TM^ 568 | Goat | 1/500 | InVitrogen (#A11036) |
| Goat anti-Mouse IgG (H+L) Highly Cross-Adsorbed Alexa Fluor™ 488 | Goat | 1/500 | InVitrogen (A11029) |
| Chicken anti-Goat IgG (H+L) Cross-adsorbed Alexa Fluor^TM^ 488 | Chicken | 1/500 | InVitrogen (#A21467) |

**Table S1 Antibodies table.**

| **Case number** | **Age** | **Sex** | **Braak NFT stage** | **Aβ phase** | **Aβ MTL** | **CERAD** |
| --- | --- | --- | --- | --- | --- | --- |
| **Patient 1 (UCL-14)** | 74 | F | 1 | 3 | 2 | 0 |
| **Patient 2 (UCL-17)** | unknown | M | 1 | 4 | 3 | 1 |
| **Patient 3 (UCL-15)** | 90 | F | 2 | 4 | 3 | 0 |
| **Patient 4 (UCL-16)** | 80 | M | 2 | / | 2 | 0 |

**Table S2 Overview of the human cohort characteristics.**
